# Human ANP32A/B are SUMOylated and utilized by avian influenza virus NS2 protein to overcome species-specific restriction

**DOI:** 10.1101/2024.07.19.604247

**Authors:** Liuke Sun, Xing Guo, Mengmeng Yu, Xue-Feng Wang, Huiling Ren, Xiaojun Wang

## Abstract

Human ANP32A/B (huANP32A/B) poorly support the polymerase activity of avian influenza viruses (AIVs), thereby limiting interspecies transmission of AIVs from birds to humans. The SUMO-interacting motif (SIM) within NS2 promotes the adaptation of AIV polymerase to huANP32A/B via a yet undisclosed mechanism. Here we show that huANP32A/B are SUMOylated by the E3 SUMO ligase PIAS2α, and deSUMOylated by SENP1. SUMO modification of huANP32A/B results in the recruitment of NS2, thereby facilitating huANP32A/B-supported AIV polymerase activity. Such a SUMO-dependent recruitment of NS2 is mediated by its association with huANP32A/B via the SIM-SUMO interaction module, where K68/K153-SUMO in huANP32A or K68/K116-SUMO in huANP32B interacts with the NS2-SIM. The SIM-SUMO interactions between huANP32A/B and NS2 function to promote AIV polymerase activity by positively regulating AIV vRNP-huANP32A/B interactions and AIV vRNP assembly. Our study offers insights into the mechanism of NS2 SIM in facilitating AIVs adaptation to mammals.

## Introduction

Influenza A viruses (IAVs) represent a significant cause of respiratory illness in both animals and humans, and pose a serious threat to human health^1^. While most influenza strains primarily circulate within avian hosts, certain highly pathogenic AIV strains (e.g., H5N1, H5N8, and H7N9) have the capacity to infect humans with potentially fatal consequences. Once these strains acquire the ability for human-to-human transmission through mutation, they may even cause influenza pandemics, presenting a substantial risk to public health. Nonetheless, the cross-species transmission of AIVs from avian hosts to mammals is limited by robust host barriers across different species^1–4^. The viral polymerase (vPol) plays a pivotal role in directing the transcription and replication of the viral genome, serving as a major determinant of the host range of IAVs^5–8^. In general, the vPol activity of AIVs is severely restricted in mammalian cells, thus hindering efficient initiation of the replication process^9,10^. Recent research has elucidated that the host factors ANP32A/B, which are members of the acidic nuclear phosphoprotein 32 kDa (ANP32) family, serve as essential cofactors for the IAV polymerase. Species-specific differences in this host factor underlie the severe restriction of AIV vPol within mammalian cells. Chicken ANP32A (chANP32A) efficiently supports AIV vPol activity owing to a specific 33 amino-acid insertion, whereas mammalian ANP32A and ANP32B lack such an insertion and are unable to effectively support AIV vPol function^11,12^. Only AIV vPol variants that have acquired adaptive mutations (e.g., PB2-E627K, and others) can proficiently utilize mammalian ANP32A/B^12–14^.

SUMOylation is a ubiquitous post-translational modification in eukaryotes that involves the covalent attachment of small ubiquitin-like modifiers (SUMO) to lysine residues on target proteins^15–17^. The mammalian SUMO family comprises four members: SUMO1, SUMO2, SUMO3 and SUMO4, with only the first three primarily participating in SUMOylation. Notably, SUMO2 shares a substantial 97% amino acid sequence similarity with SUMO3, while its similarity with SUMO1 is only 50%^15–17^. SUMO modification predominantly occurs in the nucleus and is characterized by a dynamic and reversible process. The SUMOylation mechanism encompasses SUMO activating enzymes (E1), SUMO binding enzymes (E2), and SUMO ligases (E3)^15–17^. Once covalently conjugated to a target, SUMO not only exerts direct effects on target proteins but also recruits downstream effectors through the SIM-SUMO interaction pattern. This pattern involves a non-covalent interaction between SUMO and a SUMO-interacting motif (SIM) in downstream effectors^18,19^. SIMs typically feature hydrophobic cores preceded or followed by negatively charged residues. In recent years, emerging studies have unveiled the pivotal roles of SIM-SUMO interactions in various crucial cellular processes, including liquid-liquid phase separation, histone function, DNA transcription, DNA repair, the cell cycle, and diverse host-pathogen interactions^19–21^.

Recently, our research has demonstrated that AIV NS2 possesses a conserved SIM, and AIVs utilize this SIM to help them cross the species barrier from birds to mammals by adapting their avian vPol to mammalian ANP32A/B^22^. However, the precise mechanism by which the SIM in NS2 enhances mammalian ANP32A/B-supported avian vPol activity remains unclear. Our earlier discovery^22^, revealing that the SIM in NS2 exhibits a preference for interacting with SUMO-conjugated proteins rather than with SUMO itself, led us to speculate whether NS2 relies on the SIM-SUMO interaction pattern to associate with either SUMOylated viral proteins or host factors, and thereby exerting its function in promoting mammalian ANP32A/B-supported AIV vPol activity. Here, we show that huANP32A/B are SUMOylated and thus are able to interact with NS2 via its SIM. SUMOylation of huANP32A/B is specifically fine-tuned by the E3 SUMO ligase PIAS2α and the deSUMOylation enzyme SENP1. NS2 uses its SIM to bind with SUMOylated huANP32A/B in the nucleus via a SIM-SUMO interaction pattern. This then increases huANP32A/B-supported avian vPol activity by facilitating avian vRNP assembly and vRNP-huANP32A/B interactions. We also identified that SUMOylation of huANP32A at K68/K153 or huANP32B at K68/K116 appears to be required for their association with NS2. Impairment of the SIM-SUMO mediated interactions between NS2 and huANP32A/B results in reduction of avian vPol activity and AIVs replication. Thus, SIM-SUMO interactions between viral proteins and host factors represent a new regulatory mechanism for the function of avian vPol in human cells and, therefore, also for the adaptation of AIVs.

## Results

### NS2 relies on its SIM to interact with huANP32A/B

To identify SUMOylated factors that interact with NS2 via SIM-SUMO interaction, as proposed in our previous study^22^, Flag-tagged H9N2-NS2 was co-transfected into HEK293T cells together with plasmids encoding His-SUMO1 and Myc-Ubc9. After transfection for 24 hours, cells were infected with H9N2 AIV for a further 24 hours and Flag-NS2 was purified using anti-Flag beads (Supplementary Fig. 1). Purified protein complexes were analyzed by liquid chromatography tandem mass spectrometry (LC-MS/MS), and huANP32A was identified from Flag-NS2-derived samples (Supplementary Table 1). Interestingly, a prior large-scale proteomic analysis of SUMOylated proteins identified huANP32A/B as potential SUMOylation substrates^11,23^. Consequently, we hypothesized that host factor ANP32A/B are promising candidates.

To determine whether NS2 was able to directly bind to huANP32A or huANP32B, HEK293T cells were transfected with huANP32A-Flag or huANP32B-Flag, together with GST-NS2 or GST. After 24 hours of transfection, the cells were subjected to immunoprecipitation using anti-Flag beads. As shown in Fig. 1a, GST-NS2, but not GST alone, co-precipitated with both huANP32A and huANP32B, indicating a direct interaction of NS2 with huANP32A/B. To further investigate whether the SIM of NS2 was necessary for its interaction with huANP32A/B, we utilized a SIM-defective mutant of NS2. This mutant (NS2-VE/GG) contains two glycine substitutions (V109G and E110G, VE/GG) in the core region of SIM, as described in previous reports^22^. The mutant was used for analysis of its binding with huANP32A/B in a co-IP assay. Interestingly, compared to wild-type NS2 (NS2-WT), NS2-VE/GG exhibited reduced interaction with both huANP32A and huANP32B (Fig. 1b). These results were further confirmed using bimolecular fluorescence complementation (BiFC) assays, where NS2 and huANP32A/B were expressed as fusion proteins with N- and C-terminal halves of the Venus protein, respectively. Specifically, the N-terminal residues 1 to 173 of Venus (VN) and C-terminal residues 174 to 238 (VC) were fused to the N-terminus of Myc-NS2 (VN-NS2) and to the C-terminus of the huANP32A/B-Flag (huANP32A/B-VC), respectively (Fig. 1c). When VN-NS2 and huANP32A/B-VC were expressed individually in HEK293T cells, no fluorescence signals were detected using confocal microscopy or flow cytometry. However, strong reconstituted fluorescence signals were observed when VN-NS2 was paired with huANP32A-VC or huANP32B-VC (Fig. 1d-g). Confocal microscopy analysis further revealed that the interaction between VN-NS2 and huANP32A/B-VC predominantly takes place in the nucleus (Fig. 1d). To further validate that the signal in this assay was not an artifact, the VN-NS2-VE/GG construct, bearing VE/GG mutations and showing impaired huANP32A/B binding, was co-expressed with huANP32A-VC or huANP32B-VC in HEK293T cells. As shown in Fig. 1e-g, notably faint BiFC signals were detected from the VN-NS2-VE/GG and huANP32A-VC pairs or VN-NS2-VE/GG and huANP32B-VC pairs. This confirms that the reconstituted fluorescence signals from the VN-NS2 and huANP32A/B-VC combination represent specific NS2-huANP32A/B interactions.

**Fig. 1.**
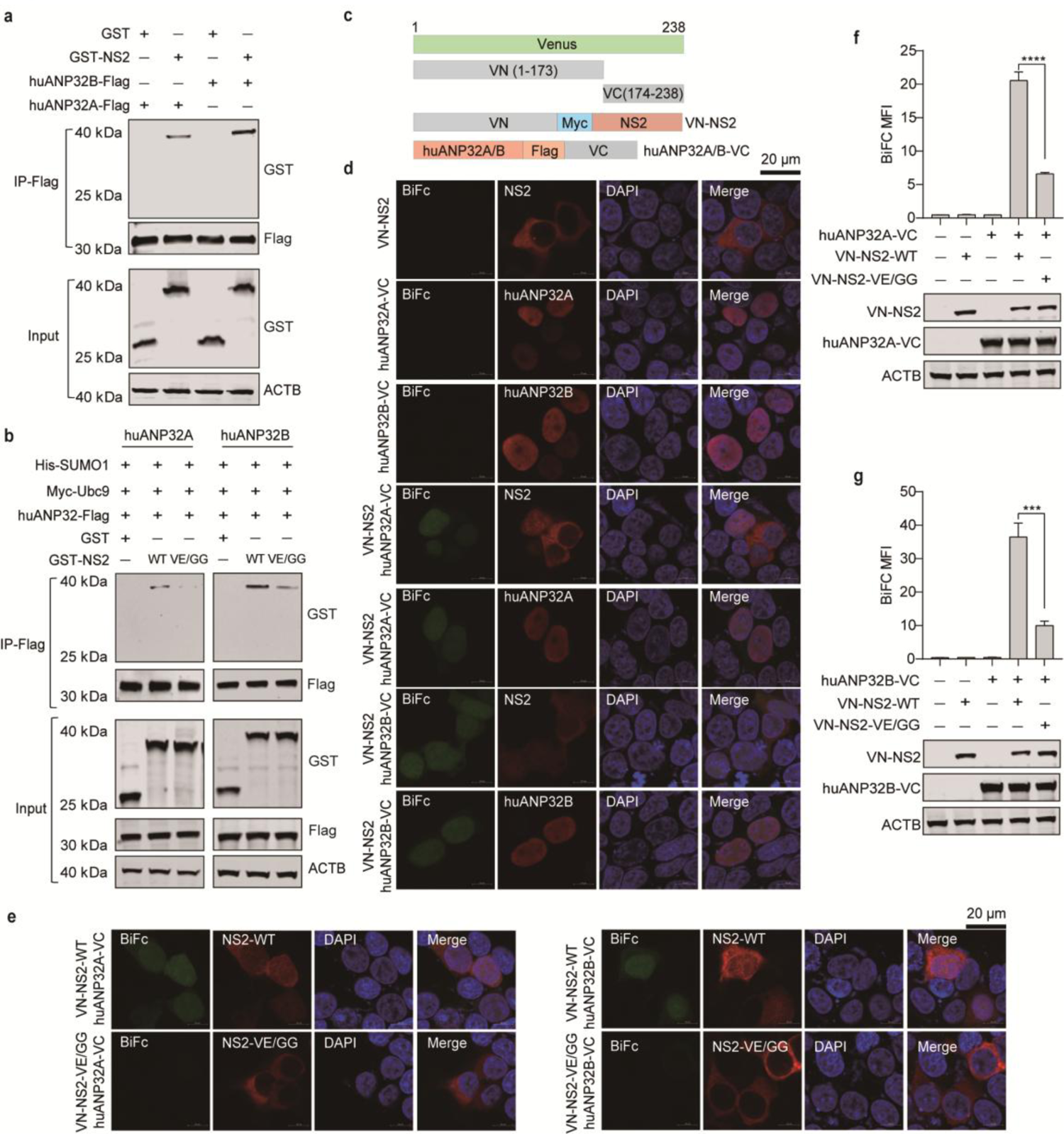
NS2 relies on its SIM to interact with huANP32A/B **a** Co-IP experiments showing the interaction of NS2 with huANP32A or huANP32B in HEK293T cells. **b** Co-IP experiments showing that mutations (V109G and E110G, VE/GG) in the SIM of NS2 suppressed the interaction between NS2 and huANP32A (left) or huANP32B (right). **c** Schematic representation of the BiFC fusion proteins. **d** Confocal experiments using a BiFC assay showing the interaction of NS2 with huANP32A or huANP32B. **e** Confocal experiments using a BiFC assay showing that VE/GG mutations suppress the interaction of NS2 with huANP32A (left) or huANP32B (right). **f, g** Flow cytometry analysis using a BiFC assay showing that NS2 interacts with huANP32A (f) or huANP32B (g) depending on its SIM. The indicated plasmids were transfected individually or in pairs into HEK293T cells. Twenty-four hours following transfection, the mean fluorescence intensity (MFI) of the BiFC fluorescence signals of the transfected cells was determined using flow cytometry. Western blots showing the protein expression of the indicated plasmids in HEK293T cells. (Error bars represent means ± SD from n = 3 independent biological replicates; \*\*\**p* < 0.001; \*\*\*\**p* < 0.0001 by unpaired Student’s t-test. In (a) and (b), experiments were independently repeated three times with consistent results.

These results collectively demonstrate that the SIM of NS2 plays a crucial role in mediating its interaction with huANP32A/B within the nucleus.

### Human ANP32A and ANP32B are SUMOylated by SUMO1, SUMO2 and SUMO3

To determine whether huANP32A/B are SUMOylated and whether this SUMOylation is influenced by AIVs infection, we initially investigated whether huANP32A and huANP32B serve as authentic SUMOylation substrates. To this end, we conducted in vivo SUMOylation assays by transfecting HEK293T cells with huANP32A-HA, Myc-Ubc9, and either His-SUMO1/2/3 or their non-conjugatable equivalents carrying G-A mutations at the two C-terminal glycine residues crucial for conjugation (His-SUMO1m/2m/3m). Following enrichment of SUMOylated proteins via histidine affinity under denaturing conditions, immunoblot analysis using an anti-HA antibody revealed three clear bands above 40 kDa for huANP32A upon overexpression of His-tagged SUMO1/2/3, but not with His-SUMO1m/2m/3m (Fig. 2a, left). Given that the molecular weights of HA-tagged huANP32A and SUMO1/2/3 are ∼33kDa and ∼15-20kDa, respectively, we speculated that huANP32A underwent modification by at least two SUMO moieties. Subsequently, we conducted in vivo SUMOylation assays with an huANP32B-HA construct, revealing that huANP32B could indeed undergo SUMOylation. The pattern of SUMOylated species is the same as that of the huANP32A, suggesting that at least two sites in huANP32B were modified by SUMOylation (Fig. 2a, right). To further substantiate these findings, we demonstrated that endogenous huANP32A/B were also SUMOylated. As shown in Fig. 2b, SUMOylated species of endogenous huANP32A and huANP32B were readily detected in HEK293T cells, but not in HEK293T-TKO cells, where the huANP32A, huANP32B and huANP32E genes were knocked out using CRISPR/Cas9 technology as described in our previous studies^24^. These results suggest that all three SUMO isoforms are covalently conjugated to huANP32A and huANP32B in HEK293T cells.

**Fig. 2.**
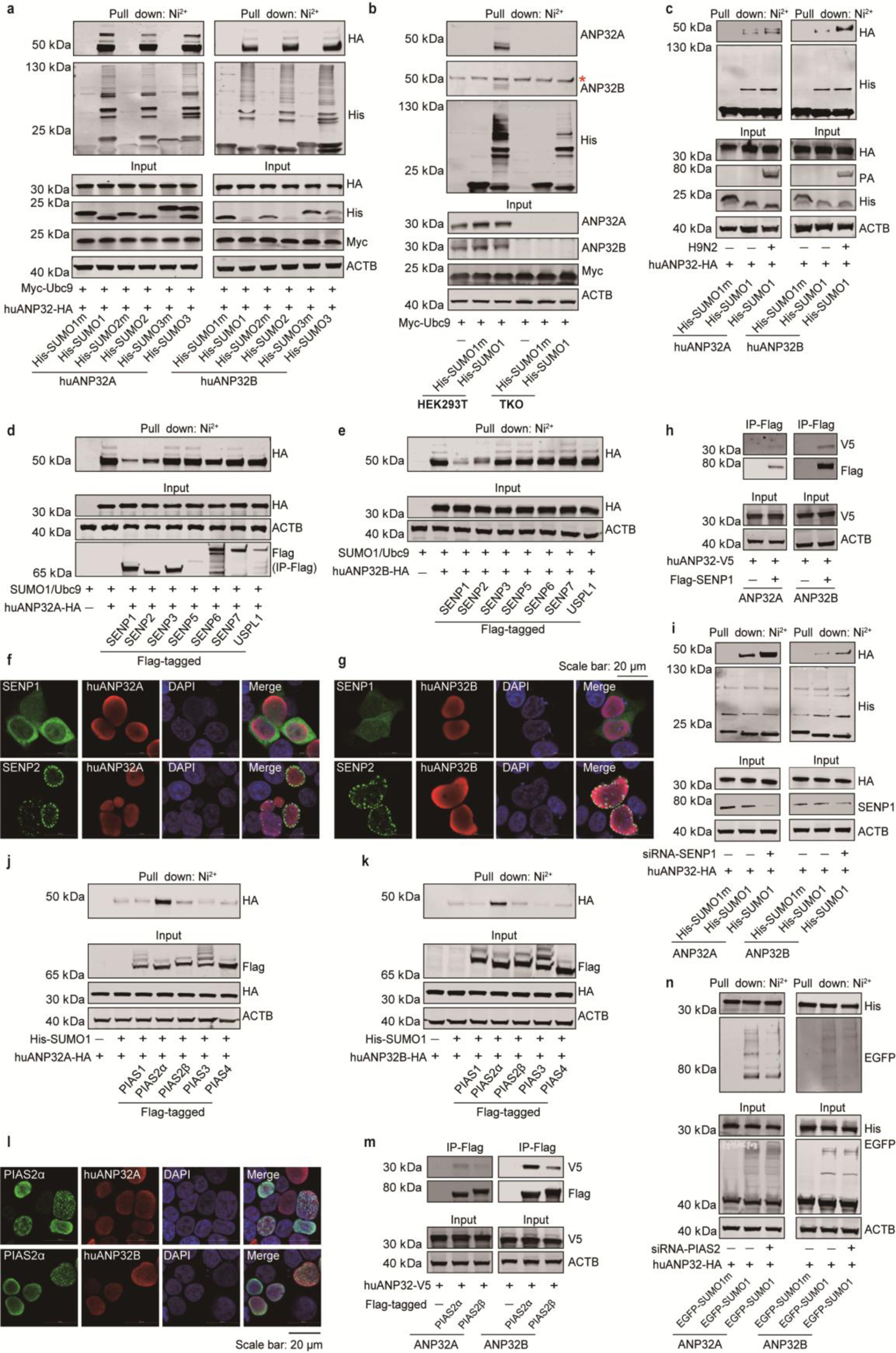
SUMOylation of huANP32A/B was regulated by the deSUMOylase SENP1 and the SUMO E3 ligase PIAS2α **a** Ni^2+^-NTA bead affinity pull-down assay results showing the modification of exogenous huANP32A (left) and huANP32B (right) with SUMO1, SUMO2, or SUMO3 in HEK293T cells. **b** Ni^2+^-NTA bead affinity pull-down assay results showing the modification of endogenous huANP32A and huANP32B with SUMO1 in HEK293T cells. *indicates non-specific protein band. **c** Ni^2+^-NTA bead affinity pull-down assay results showing that the SUMOylation of huANP32A (left) or huANP32B (right) in HEK293T cells was notably enhanced by H9N2 AIV infection. HEK293T cells were transfected with the indicated plasmids for 24 hours followed by H9N2 AIV infection (MOI = 0.1) for another 24 hours. Cells were then harvested for in vivo SUMOylation assays and immunoblotting analysis. **d, e** Ni^2+^-NTA bead affinity pull-down assay results showing that the SUMOylation of huANP32A (d) or huANP32B (e) in HEK293T cells was notably reduced by overexpression of SENP1 or SENP2. **f, g** Co-localization of huANP32A-V5 (f) or huANP32B-V5 (g) with Flag-SENP1 and Flag-SENP2 was analyzed using immunofluorescence staining. Scale bar: 20 μm. **h** Co-IP experiments showing the interaction of SENP1 with huANP32A (left) or huANP32B (right) in HEK293T cells. **i** Ni^2+^-NTA bead affinity pull-down assay results showing that knockdown of endogenous SENP1 enhanced the SUMOylation of both huANP32A and huANP32B in HEK293T cells. **j, k** Ni^2+^-NTA bead affinity pull-down assay results showing that the SUMOylation of huANP32A (J) or huANP32B (K) in HEK293T cells was notably enhanced by overexpression of PIAS2α. **l** Co-localization of huANP32A-V5 or huANP32B-V5 with Flag-PIAS2α was analyzed using immunofluorescence staining. Scale bar: 20 μm. **m** Co-IP experiments showing the interaction of PIAS2α with huANP32A (left) or huANP32B (right) in HEK293T cells. **n** Ni^2+^-NTA bead affinity pull-down assay results showing that knockdown of endogenous PIAS2 reduced the SUMOylation of both huANP32A and huANP32B in HEK293T cells. In (a to e), (h to k), (m) and (n), experiments were independently repeated three times with consistent results.

Previous studies have demonstrated that IAVs infection contributes to a global increase in the level of SUMOylation in host cells^23^. To examine whether AIVs infection specifically enhances the SUMOylation levels of huANP32A and huANP32B, HEK293T cells were transfected with HA-tagged huANP32A or huANP32B along with His-SUMO1 or its non-conjugatable form (His-SUMO1m). Following a 24-hour transfection period, the cells were subsequently infected with or without H9N2 AIV for an additional 24 hours. The SUMOylation of huANP32A/B was then assessed through in vivo SUMOylation assays. As shown in Fig. 2c, the SUMOylation of both huANP32A and huANP32B exhibited an increase upon H9N2 AIV infection.

Collectively, these findings conclusively establish that both huANP32A and huANP32B act as targets for SUMOylation, and that their SUMOylation modifications are positively regulated by AIVs infection.

### SUMOylation of huANP32A/B was controlled by the deSUMOylating enzymes SENP1 and E3 SUMO ligase PIAS2α

SUMOylation represents a dynamic and reversible cellular process governed by deSUMOylating enzymes, and sentrin-specific proteases (SENPs) are the major deSUMOylating enzymes in mammalian cells^25^. To pinpoint the specific SUMO proteases responsible for the deSUMOylation of huANP32A/B, we co-transfected all known human nuclear SUMO-specific proteases (SENP1-3, SENP5-7 and USPL1) with huANP32A-HA or huANP32B-HA, along with Myc-Ubc9 and His-SUMO1, into HEK293T cells. Levels of huANP32A/B SUMOylation were then assessed through in vivo SUMOylation assays. We observed that co-expression of SENP1 and SENP2 notably decreased huANP32A/B SUMOylation (Fig. 2d, e). Immunofluorescence staining revealed that SENP1, but not SENP2, co-localized with huAN32A or huANP32B in the nucleus (Fig. 2f, g). Consequently, we focused on addressing the role of SENP1 in huANP32A/B deSUMOylation. To substantiate this role, we investigated the interaction between SENP1 and huANP32A or huANP32B in HEK293T cells. As shown in Fig. 2h, SENP1 specifically interacts with huANP32A and huANP32B. Most importantly, depletion of endogenous SENP1 led to a considerable enhancement of huANP32A/B SUMOylation (Fig. 2i). These findings indicate that SENP1 targets huANP32A/B for deSUMOylation.

The primary SUMO E3 ligases in mammalian cells belong to the PIAS family proteins^26^. The human PIAS gene family is known to comprise four members: PIAS1, PIASx (also known as PIAS2), PIAS3 and PIASy (also known as PIAS4), with PIAS2 existing in two isoforms (PIAS2α and PIAS2β). To determine the putative E3 SUMO ligase for huANP32A/B SUMOylation, His-SUMO1 and each of the five PIAS(s) were co-transfected with huANP32A-HA or huANP32B-HA into HEK293T cells, and huANP32A/B SUMOylation was assessed by performing in vivo SUMOylation assays. Of the tested isoforms, PIAS2α prominently enhanced huANP32A/B SUMOylation, while the other PIAS variants did not exhibit any such effects (Fig. 2j, k). Additionally, PIAS2α specifically colocalized with huANP32A and huANP32B within the nucleus (Fig. 2l). We confirmed the specific interaction between PIAS2α and huANP32A as well as huANP32B using a co-IP assay (Fig. 2m). Depletion of endogenous PIAS2 resulted in defects in huANP32A/B SUMOylation (Fig. 2n; Supplementary Fig. 2). These results clearly indicate that PIAS2α plays a crucial role in directing huANP32A/B for SUMOylation.

These findings collectively establish that both huANP32A and huANP32B undergo SUMOylation catalyzed by the E3 ligase PIAS2 and subsequent deSUMOylation mediated by SENP1.

### NS2 engages with huANP32A/B via a SIM-SUMO-dependent manner

To explore the impact of huANP32A/B SUMOylation on their interactions with NS2, we investigated whether the huANP32A/B-NS2 interactions were affected by the alterations in huANP32A/B SUMOylation levels. This was achieved by overexpressing the single SUMO E2 conjugating enzyme (Ubc9), employing the targeted deSUMOylation enzyme SENP1 for huANP2A/B SUMOylation, or by administering a selective inhibitor of the SUMOylation enzymatic cascade (TAK-981). As shown in Fig. 3a, co-expression of Ubc9 led to a substantial increase in huANP32A/B SUMOylation. Consequently, Ubc9 heightened the huANP32A/B-NS2 interactions (Fig. 3b, c). Conversely, co-expression of SENP1 diminished huANP32A/B SUMOylation, thereby inhibiting the interaction of huANP32A/B with NS2 (Fig. 3d, e). Moreover, TAK-981 treatment not only impeded huANP32A SUMOylation but also disrupted the huANP32A-NS2 interaction (Fig. 3f, g). These findings unequivocally underscore the crucial role of huANP32A/B SUMOylation in facilitating their association with NS2.

**Fig. 3.**
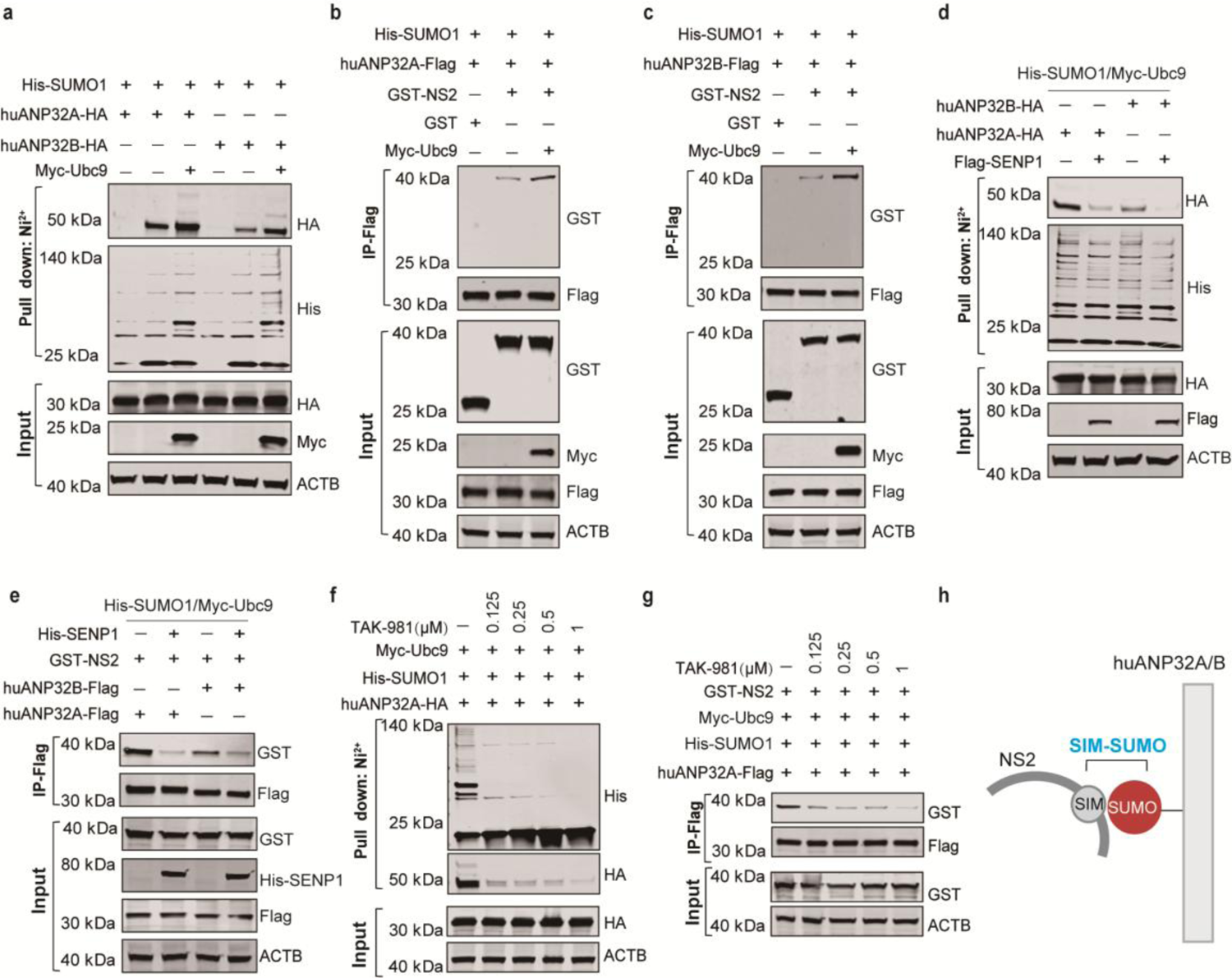
SUMOylation of huANP32A and huANP32B determines their association with NS2 **a** Ni^2+^-NTA bead affinity pull-down assay results showing that SUMOylation of huANP32A or huANP32B in HEK293T cells was enhanced by overexpression of Ubc9. **b**, **c** Co-IP experiments showing that overexpression of Ubc9 promoted the interaction between NS2 and huANP32A (b) or huANP32B (c). **d** Ni^2+^-NTA bead affinity pull-down assay results showing that SUMOylation of huANP32A or huANP32B in HEK293T cells was reduced by overexpression of SENP1. **e** Co-IP experiments showing overexpression of SENP1 suppressed the interaction between NS2 and huANP32A or huANP32B. **f** Ni^2+^-NTA bead affinity pull-down assay results showing the SUMOylation of huANP32A in HEK293T cells was reduced by treatment with TAK-981. **g** Co-IP experiments showing that treatment with TAK-981 suppressed NS2-huANP32A interaction. **h** Illustration of the interaction model between NS2 and huANP32A/B mediated by the SIM-SUMO working pattern. In (a to g), experiments were independently repeated at least twice with consistent results.

Collectively, the above results demonstrate that the interactions between huANP32A/B and NS2 relies on the canonical SIM-SUMO interaction module, where SUMO attached to huANP32A/B interacts with the SIM on NS2 (Fig. 3h).

### SIM-SUMO interactions between huANP32A/B and NS2 are required for the avian polymerase-enhancing function of NS2

We next intended to assess the necessity of SIM-SUMO interactions between huANP32A/B and NS2 for the avian polymerase-enhancing function of NS2. To this end, we proceeded to directly assess whether disruption of these interactions, achieved through either the destabilization of NS2-SIM integrity or inhibition of huANP32A/B SUMOylation, would affect avian vPol activity when supported by huANP32A/B in the presence of NS2 (Fig. 4a). As shown in Fig. 4b, c, disruption of NS2-SIM integrity caused NS2 to lose its avian vPol-enhancing properties in HEK293T-TKO cells reconstituted with either huANP32A or huANP32B, which is consistent with our previous study^22^. These results support the hypothesis that the SIM-SUMO mediated interactions between NS2 and huANP32A/B are indispensable for the ability of NS2 to enhance huANP32A/B-supported avian vPol activity.

**Fig. 4.**
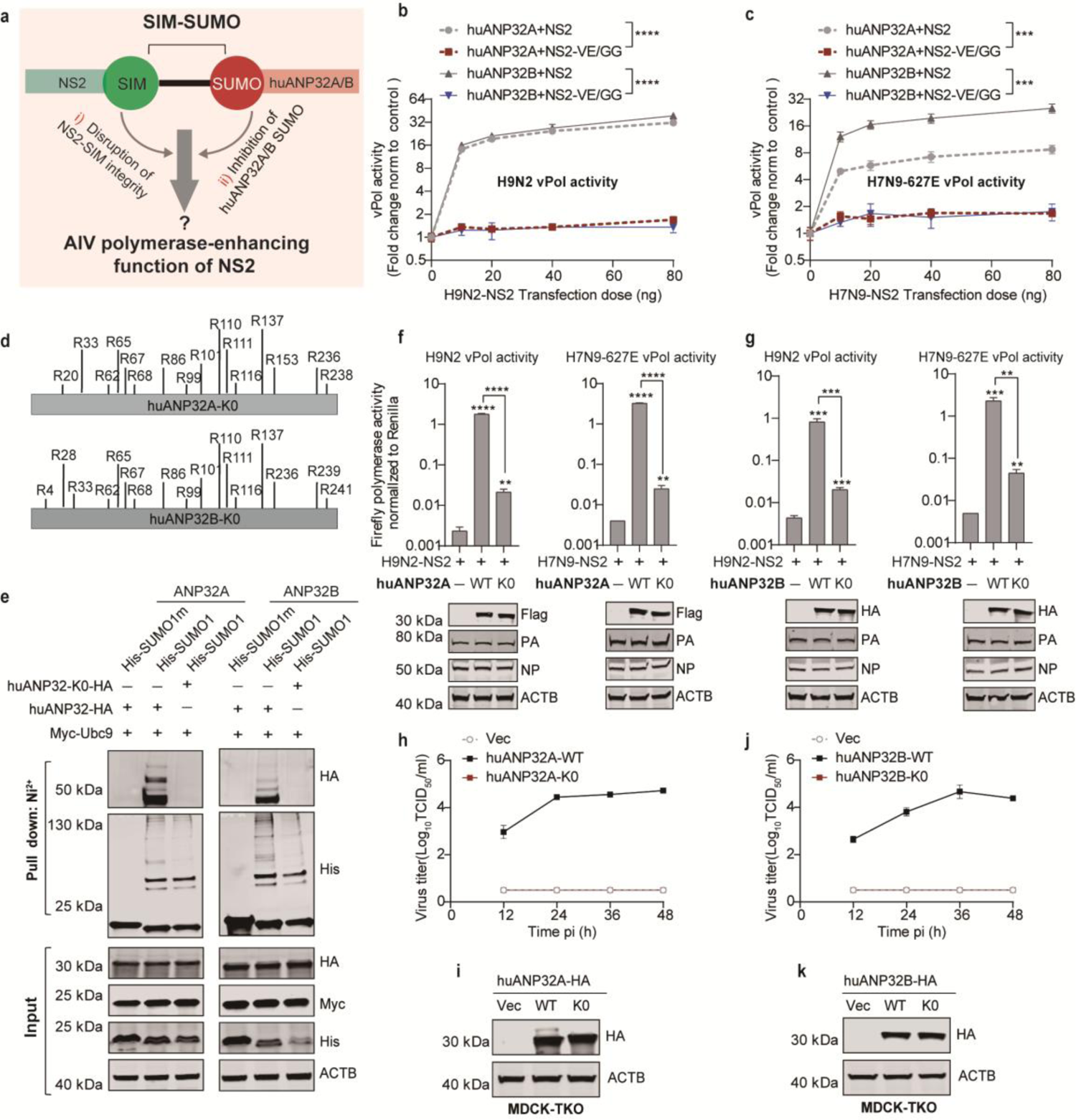
The function of NS2 in supporting avian vPol activity depends on its interaction with huANP32A/B via the SIM-SUMO working pattern **a** Schematic illustrating two strategies employed for the investigation of whether the SIM-SUMO interaction between NS2 and huANP32A/B is required for NS2 to promote huANP32A/B-supported AIV vPol activity. **b, c** Minigenome assays in HEK293T-TKO cells showing that disruption of H9N2 or H7N9 NS2-SIM integrity impairs the ability of NS2 to support huANP32A/B-supported H9N2 or H7N9 (PB2-627E) vPol activity, respectively. **d** Schematic model of the generation of lysine-free mutant of huANP32A/B (huANP32A/B-K0). **e** Ni^2+^-NTA bead affinity pull-down assay results showing that huANP32A/B-K0 could not be SUMOylated in HEK293T cells. **f, g** Minigenome assays in HEK293T-TKO cells showing that huANP32A-K0 (f) or huANP32B-K0 (g) supported H9N2 or H7N9 (PB2-627E) vPol activity was greatly reduced in the presence of NS2. The accompanying Western blots show the expression of ANP32A-Flag/ANP32B-HA constructs and vRNP components (PA/NP). **h** Replication kinetics of avian H9N2 virus. MDCK-TKO cells or those tranfected with huANP32A-HA or huANP32A-K0-HA were infected (MOI = 0.01), and viral titers were determined at the indicated time points. **i** Western blots showing equal expression of HA-tagged ANP32A in transfected MDCK-TKO cells. **j** The replication kinetics of avian H9N2 virus were assessed in MDCK-TKO cells transfected with either empty vector, huANP32B-HA or huANP32B-K0-HA (MOI = 0.01). Viral titers were determined at the indicated time points. **k** Western blots showing equal expression of HA-tagged ANP32B in transfected MDCK-TKO cells. In (b), (c), (f to h) and (j), error bars represent mean ± SD from n = 3 independent biological replicates; \*\**p* < 0.01, \*\*\**p* < 0.001, and \*\*\*\**p* < 0.0001 by two-way ANOVA (b and c) or unpaired Student’s t-test (f and g).

We then proceeded to investigate the impact of inhibiting SIM-SUMO interactions on the activity of huANP32A/B-supported avian vPol by reducing the level of huANP32A/B SUMOylation. To achieve this, we engineered a SUMOylation-deficient mutant of huANP32A, denoted as huANP32A-K0, with lysine-to-arginine substitutions at all sixteen lysine residues of the protein. Similarly, a counterpart mutant of huANP32B, termed huANP32B-K0, was generated with lysine-to-arginine substitutions at all seventeen lysine residues (Fig. 4d). Theoretically, huANP32A/B-K0 should lose its capability to be SUMOylated and would therefore be unable to engage in the SIM-SUMO interaction mode crucial for interacting with NS2. As we expected, the SUMOylation assays we conducted in HEK293T cells confirmed that neither huANP32A-K0 nor huANP32B-K0 were able to undergo SUMOylation (Fig. 4e). Subsequent polymerase activity assays performed in TKO cells revealed a significant reduction in the ability of NS2 to enhance H9N2 and H7N9 (PB2-627E) vPol activity when supported by huANP32A/B-K0 compared to wild-type huANP32A/B (Fig. 4f, g).

To evaluate whether the observed changes in polymerase activity correlate with viral replication, we firstly established MDCK-TKO cells by knocking out the triple canine ANP32A, ANP32B, and ANP32E genes using CRISPER-Cas 9 technology (Supplementary Fig. 3). Multi-cycle replication assays of avian H9N2 virus were then subsequently conducted in MDCK-TKO cells reconstituted with empty vector, huANP32A-WT or huANP32A-K0, respectively. Notably, virus growth kinetics revealed that huANP32A-K0 has lost its ability to support the replication of avian H9N2 viruses (Fig. 4h). Western blot analysis confirmed the equal expression of huANP32A proteins in these transfected cells (Fig. 4i). We performed similar experiments for huANP32B and huANP32B-K0 and obtained similar observations that huANP32B-K0 failed to support the replication of avian H9N2 viruses (Fig. 4j, k).

Taken together, these results demonstrate that disruption of either huANP32A/B SUMO or NS2 SIM sites attenuates their interaction and impairs huANP32A/B-supported AIV polymerase activity. This highlights the crucial role of SIM-SUMO interactions between huANP32A/B and NS2 in facilitating NS2 function in promoting AIV polymerase activity, especially when supported by huANP32A/B.

### SUMOylation of huANP32A at K68/K153 determines its association with NS2

Human ANP32A comprises sixteen lysine residues. Our objective was to pinpoint the primary site for huANP32A SUMOylation crucial for its association with NS2. Given that the SIM-SUMO interactions between NS2 and huANP32A/B are required for NS2 to promote huANP32A/B-supported avian vPol activity, it is reasonable to expect that reintroducing into huANP32A-K0 the major SUMOylation site present in wild-type huANP32A, which determines its association with NS2, will enhance the support of avian vPol activity by huANP32A-K0 in the presence of NS2. To explore this, a series of huANP32A-K0 mutants were generated, in which each of the sixteen lysines present in wild-type huANP32A was individually mutated back (R-K) into the huANP32A-K0 construct. Western blot analysis confirmed comparable expression levels of all indicated constructs (Supplementary Fig. 4). Subsequently, these huANP32A-K0 constructs were assessed for their ability to support avian vPol activity by performing polymerase activity assays in TKO cells. The results revealed that in the presence of NS2, four mutants, huANP32A-K0-R68K, huANP32A-K0-R116K, huANP32A-K0-R153K and huANP32A-K0-R236K, supported H9N2 vPol activity to a greater extent than huANP32A-K0, indicating that SUMOylation at sites K68, K153, K116 and K236 is crucial for huANP32A support of avian vPol activity (Fig. 5a). In comparison to the other three mutations, the R236K mutation exhibited the weakest impact on polymerase activity, resulting in less than a 1-fold effect. Consequently, the R236K mutation was excluded from subsequent experiments. Double, and triple R-K mutations at these sites were then further introduced into huANP32A-K0 constructs (Fig. 5b, c), and these modified constructs were individually reconstituted into TKO cells for polymerase activity assays. As shown in Fig. 5d, e, double (R68K/R153K) and triple (R68K/R116K/R153K) point mutants supported significantly higher levels of H9N2 and H7N9 vPol activity compared to single point mutants, including R68K and R153K. However, the introduction of the R116K mutation into double mutants (R68K/R153K) did not further enhance its ability to support avian vPol activity, suggesting that K68 and K153 are the two major sites for SUMOylation in huANP32A that determine its association with NS2. Additionally, by establishing two different MDCK-TKO cell lines stably expressing either Flag-tagged huANP32A-K0 or huANP32A-K0-R68K/R153K, we observed that avian H9N2 viruses failed to establish a productive infection in either control MDCK-TKO cells or MDCK-TKO cells expressing Flag-tagged huANP32A-K0. However, MDCK-TKO cells expressing Flag-tagged huANP32A-K0-R68K/R153K efficiently supported the replication of avian H9N2 viruses (Fig. 5f, g).

**Fig. 5.**
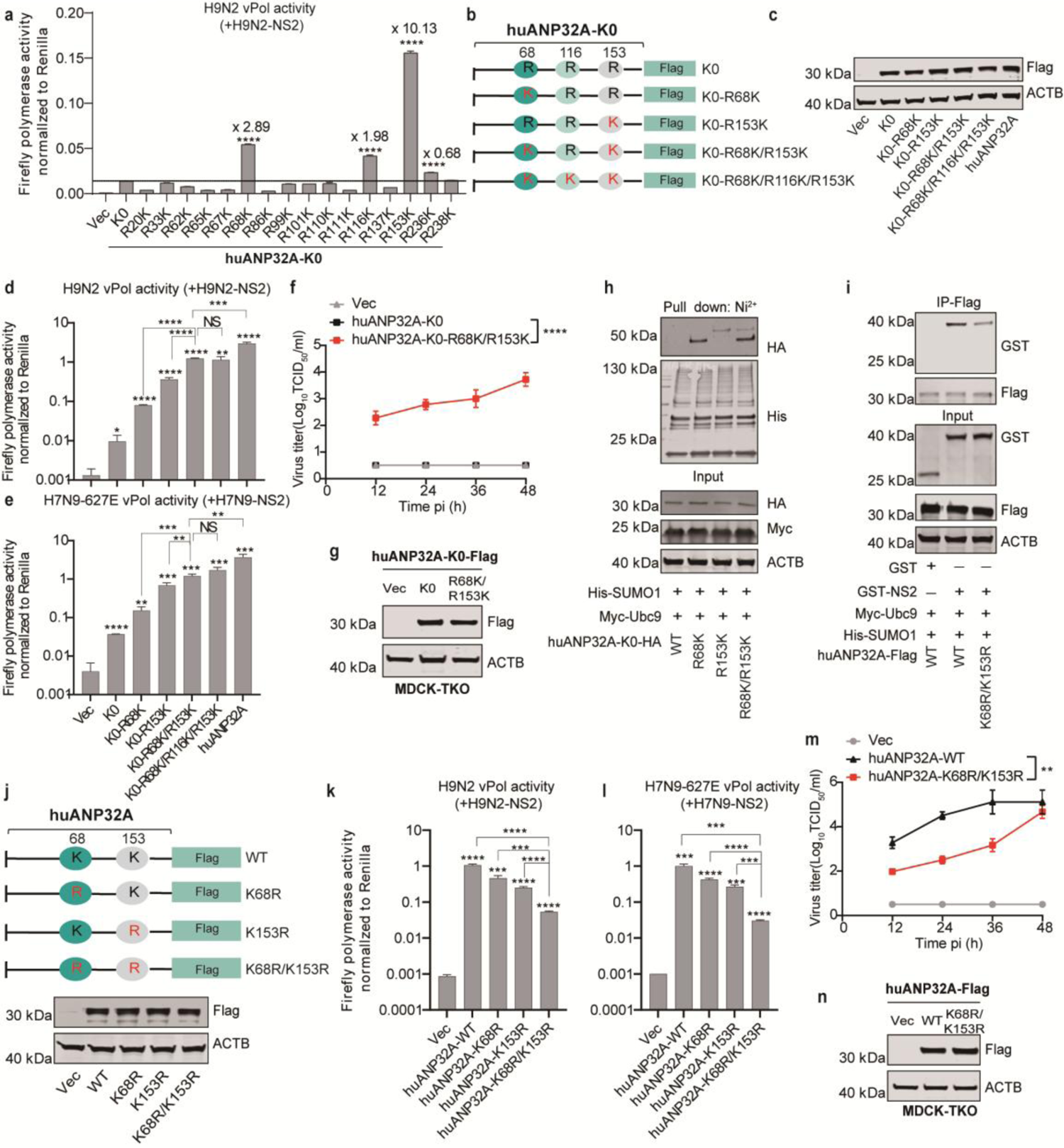
The SIM of NS2 binds to the SUMO at the K68/K153 sites of huANP32A. **a** Minigenome assays in HEK293T-TKO cells comparing the effect of the indicated constructs on H9N2 vPol activity. Values above K0 were statistically analyzed using one-way ANOVA followed by a Dunnett’s multiple comparisons test against huANP32A-K0 (n = 4 independent biological replicates; \*\*\*\**p* < 0.0001). **b** The huANP32A-K0 mutants generated. **c** Western blots demonstrate comparable expression levels for all indicated constructs. **d, e** Minigenome assays in HEK293T-TKO cells comparing the effect of the indicated huANP32A-K0 constructs on the H9N2 (D) and H7N9 (PB2-627E) (E) vPol activity. **f** The replication kinetics of the avian H9N2 virus were evaluated in control MDCK-TKO cells or those stably expressing huANP32A-K0-Flag or its mutant (MOI = 0.01), with viral titers determined at the indicated time points. **g** Western blot analysis of MDCK-TKO cells stably reconstituted with the indicated huANP32A-K0-Flag constructs or empty vector. **h** Ni^2+^-NTA bead affinity pull-down assay showing that the K68 and K153 sites of huANP32A can be modified by SUMO1. **i** Co-IP experiments showing that K68R/K153R mutations suppress NS2-huANP32A interaction. **j** The huANP32A mutants generated, and western blots demonstrate comparable expression levels for all indicated constructs. **k**, **l** Minigenome assays in HEK293T-TKO cells comparing the effect of the indicated huANP32A constructs on the H9N2 (k) and H7N9 (PB2-627E) (l) vPol activity. **m** The replication kinetics of the avian H9N2 virus were evaluated in control MDCK-TKO cells or those stably expressing huANP32A-Flag or huANP32A-K68R/K153R-Flag (MOI = 0.01), with viral titers determined at the indicated time points. **n** Western blot analysis of control MDCK-TKO cells or those stably expressing indicated huANP32A-Flag constructs. In (d to f) and (k to m), error bars represent the mean ± SD of n = 3 independent biological replicates; NS, not significant; \**p* < 0.05; \*\**p* < 0.01; \*\*\**p* < 0.001; \*\*\*\**p* < 0.0001 by unpaired Student’s t-test (d, e, k and l) or two-way ANOVA (f and m). In (h and i), experiments were independently repeated at least twice with consistent results.

To ascertain whether the K68 and K153 sites in huANP32A function as SUMOylation sites that determine its association with NS2, we first used in vivo SUMOylation assays to demonstrate that SUMOylation occurs at K68 and K153 in huANP32A. To this end, HEK293T cells were co-transfected with His-SUMO1, Myc-Ubc9, huANP32A-K0-HA and its different mutants. Twenty-four hours following initial transfection, cell lysates were prepared for SUMOylation assays. As shown in Fig. 5h, a signal band above 40 kDa was observed for huANP32A-K0-R68K or huANP32A-K0-R153K, indicating that K68 and K153 are indeed SUMOylation sites for huANP32A. However, the reason for the slower migration of huANP32A-K0-R153K-SUMO compared to huANP32A-K0-R68K-SUMO remains unclear and requires further investigation.

To further validate the crucial role of K68/K153-mediated SUMOylation in the SIM-SUMO interactions between huANP32A/B and NS2, we performed a co-IP assay. As shown in Fig. 5i, huANP32A-K68R/K153R co-precipitated a considerably smaller amount of NS2 than did wild type huANP32A, suggesting that SUMOylation of huANP32A at K68/K153 indeed determines its association with NS2. Interestingly, the K68R/K153R mutations did not affect the overall level of huANP32A SUMOylation (Supplementary Fig. 5a). This phenomenon implies that there are compensatory effects for huANP32A SUMOylation when introducing the K68R and K153R mutations, and only site-specific SUMOylation in huANP32A could be recognized by the SIM in NS2. Importantly, sequence alignments revealed that the K68 and K153 sites are well-conserved among ANP32A proteins from other mammalian species (Supplementary Fig. 6). Immunofluorescence staining revealed that both huANP32A-WT and huANP32A-K68R/K153R mutant were predominantly localized in the nucleus (Supplementary Fig. 7a).

In line with the impairment observed in the huANP32A-NS2 interaction due to the K68R/K153R mutations, these two mutations significantly diminished the ability of huANP32A to support H9N2 and H7N9 (PB2-627E) vPol activity (Fig. 5j-L), as well as H9N2 AIV replication (Fig. 5m, n).

Taken together, these findings strongly suggest that SUMOylation at K68/K153 of huANP32A mediates its interaction with the SIM in NS2, which is required for NS2 to promote avian vPol activity and AIV replication.

### SUMOylation of huANP32B at K68/K116 determines its association with NS2

To pinpoint the essential sites of SUMOylation in huANP32B crucial for its association with NS2, we employed the same experimental approach as for huANP32A. As shown in Fig. 6a and Supplementary Fig. 8, both the R68K and R116K mutations markedly enhanced the ability of huANP32B-K0 to support H9N2 vPol activity in the presence of NS2. Subsequently, we found that the double point mutant (R68K/R116K) supported H9N2 and H7N9 (PB2-627E) vPol activity to a significant extent when compared to single point mutants, including R68K and R116K (Fig. 6b-e), Additionally, these alterations in polymerase activity were mirrored in viral replication, as the introduction of the R68K/R116K mutation into huANP32B-K0 reinstated its ability to support the replication of H9N2 AIV (Fig. 6f, g).

**Fig. 6.**
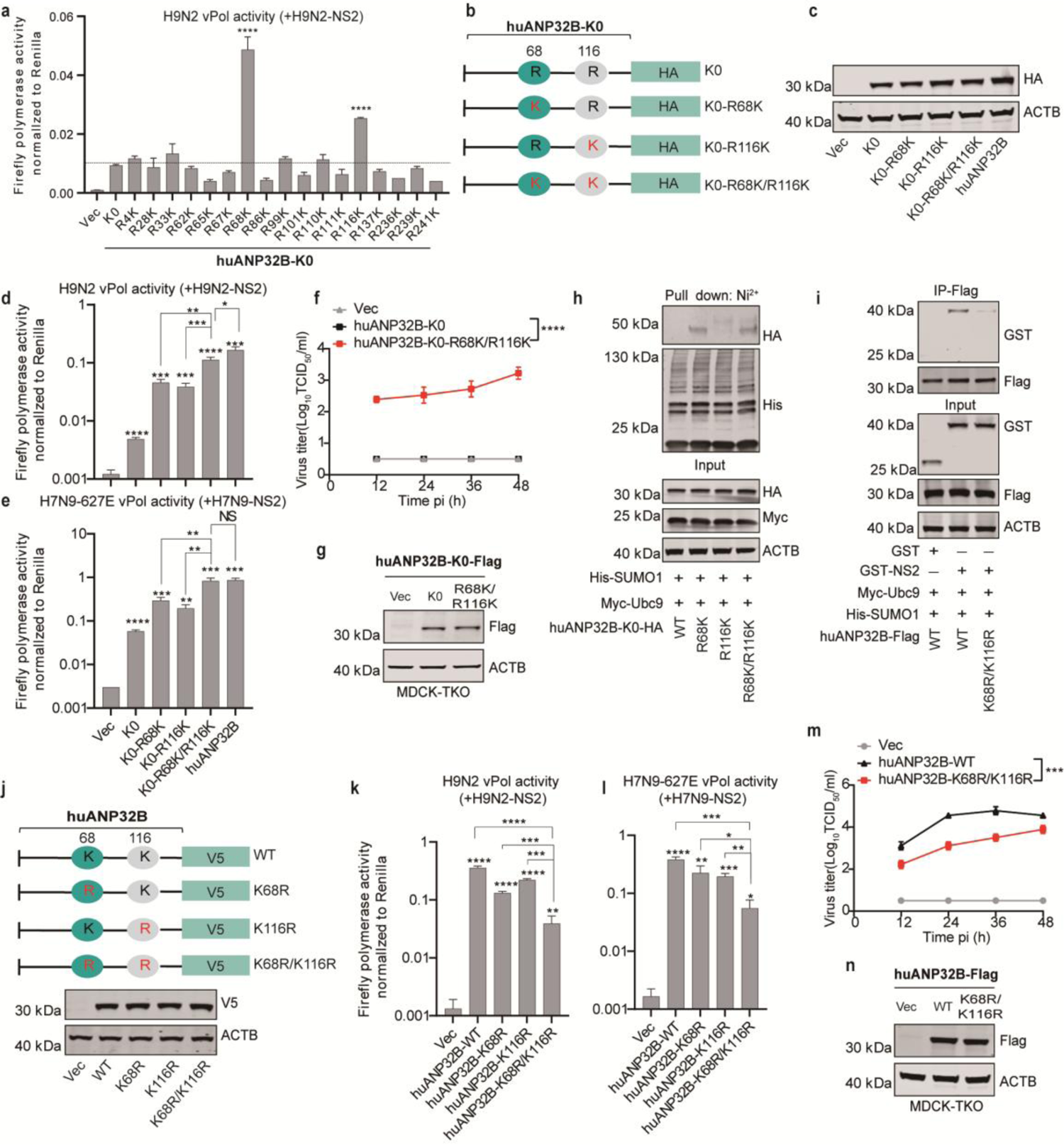
The SIM of NS2 binds to the SUMO at the K68/K116 sites of huANP32B. **a** Minigenome assays in HEK293T-TKO cells comparing the effect of indicated huANP32B-K0 constructs on the H9N2 vPol activity. Values higher than K0 were selected for statistical analysis using one-way ANOVA followed by a Dunnett‘s multiple comparisons test against huANP32B-K0 (n = 3 independent biological replicates; \*\*\*\**p* < 0.0001). **b** The huANP32B-K0 mutants generated. **c** Western blots showing that all indicated constructs of huANP32B are expressed at comparable levels. **d**, **e** Minigenome assays in HEK293T-TKO cells comparing the effect of the indicated huANP32B-K0 constructs on the H9N2 (d) and H7N9 (PB2-627E) (e) vPol activity in the presence of NS2. **f** The replication kinetics of the avian H9N2 virus were evaluated in control MDCK-TKO cells or those stably expressing huANP32B-K0-Flag or its mutant (MOI = 0.01), with viral titers determined at the indicated time points. **g** Western blot analysis of control MDCK-TKO cells or those stably expressing indicated huANP32B-K0-Flag constructs. **h** Ni^2+^-NTA bead affinity pull-down assay showing that the K68 and K116 sites of huANP32B can be modified by SUMO1. **i** Co-IP experiments showing that K68R/K116R mutations suppress NS2-huANP32B interaction. **j** The huANP32B mutants generated, and western blots showing that all indicated constructs were expressed at comparable levels. **k, l** Minigenome assays in HEK293T-TKO cells comparing the effect of the indicated huANP32B constructs on the H9N2 (k) and H7N9 (PB2-627E) (l) vPol activity. **m** The replication kinetics of the avian H9N2 virus were evaluated in control MDCK-TKO cells or those stably expressing huANP32B-Flag or its mutant (MOI = 0.01), with viral titers determined at the indicated time points. **n** Western blot analysis of control MDCK-TKO cells or those stably expressing indicated huANP32B-Flag constructs. In (d to f) and (k to m), error bars represent the mean ± SD of n = 3 independent biological replicates; NS, not significant; \**p* < 0.05; \*\**p* < 0.01; \*\*\**p* < 0.001; \*\*\*\**p* < 0.0001 by unpaired Student’s t-test (d, e, k and l) or two-way ANOVA (f and m). In (h and i), experiments were independently repeated at least twice with consistent results.

To confirm that K68 and K116 serve as the sites for SUMOylation in huANP32B, we conducted SUMOylation assays in HEK293T cells by co-transfecting His-SUMO1 with Myc-Ubc9, huANP32B-K0-HA and its various mutants. Following transfection for 24 hours, cell lysates were prepared for SUMOylation assays. As shown in Fig. 6h, a signal band above 40 kDa was observed for huANP32B-K0-R68K or huANP32B-K0-R116K, indicating that K68 and K116 are indeed SUMOylation sites for huANP32B. Similar to huANP32A-K0-R153K-SUMO, the migration of huANP32B-K0-R116K-SUMO was slower than that of huANP32B-K0-R68K-SUMO. Additionally, a co-IP assay was conducted, revealing that K68R/K116R mutations impede the NS2-huANP32B interaction (Fig. 6i). Intriguingly, the K68R/K116R mutations also did not affect the overall SUMOylation level of huANP32B (Supplementary Fig. 5b). These findings suggest that there are compensatory effects on huANP32B SUMOylation upon the introduction of K68R and K116R mutations, further supporting the notion that site-specific SUMOylation of huANP32B mediates its interaction with the SIM in NS2. Interestingly, the K68 and K116 sites are well-conserved among ANP32B proteins from other mammalian species (Supplementary Fig. 6). Immunofluorescence staining indicated that both huANP32B-WT and huANP32B-K68R/K116R mutant predominantly localized in the nucleus (Supplementary Fig. 7b).

In line with the impairment observed in the huANP32B-NS2 interaction due to the K68R/K116R mutation, these two mutations significantly diminished the ability of huANP32B to support H9N2 and H7N9 (PB2-627E) vPol activity (Fig. 6j-l), as well as H9N2 AIV replication (Fig. 6m, n).

Taken together, these data suggest that SUMOylation of huANP32B at the K68/K116 sites is required for the SIM-SUMO interaction between NS2 and huANP32B and that impairment of this interaction results in reduced avian vPol activity and AIV replication.

### SIM-SUMO interactions between NS2 and huANP32A/B function to promote avian vPol activity by positively regulating AIV vRNP-huANP32A/B interactions and AIV vRNP assembly

Based on the aforementioned findings, it is evident that the SIM-SUMO interactions between NS2 and huANP32A/B are indispensable for NS2 to effectively enhance avian vPol activity in human cells. Our previous work has demonstrated that NS2-SIM functions in promoting avian vPol activity by positively regulating vRNP-huANP32A/B interactions and avian vRNP assembly^22^. Building on this, we aimed to further investigate this phenomenon by disrupting the SUMOylation of huANP32A/B, which is crucial for their association with NS2.

We first assessed the impact of huANP32A SUMOylation on avian vRNP-huANP32A/B interactions in TKO cells reconstituted with the indicated huANP32A-Flag constructs and plasmids encoding H9N2 vRNP. As shown in Fig. 7a, NS2 enhanced the H9N2 vRNP-huANP32A interaction, but failed to enhance H9N2 vRNP-huANP32A-K0 interaction (Fig. 7a, b). However, NS2 effectively promoted the interaction between huANP32A-K0-R68K/R153K and vRNP (Fig. 7b). Furthermore, we observed that K68R/K153R mutations impaired the huANP32A-vRNP interaction (Fig. 7c). For huANP32B, we performed similar experiments with the H9N2 vRNP and the indicated huANP32B constructs and obtained similar results (Fig. 7d-f).

**Fig. 7.**
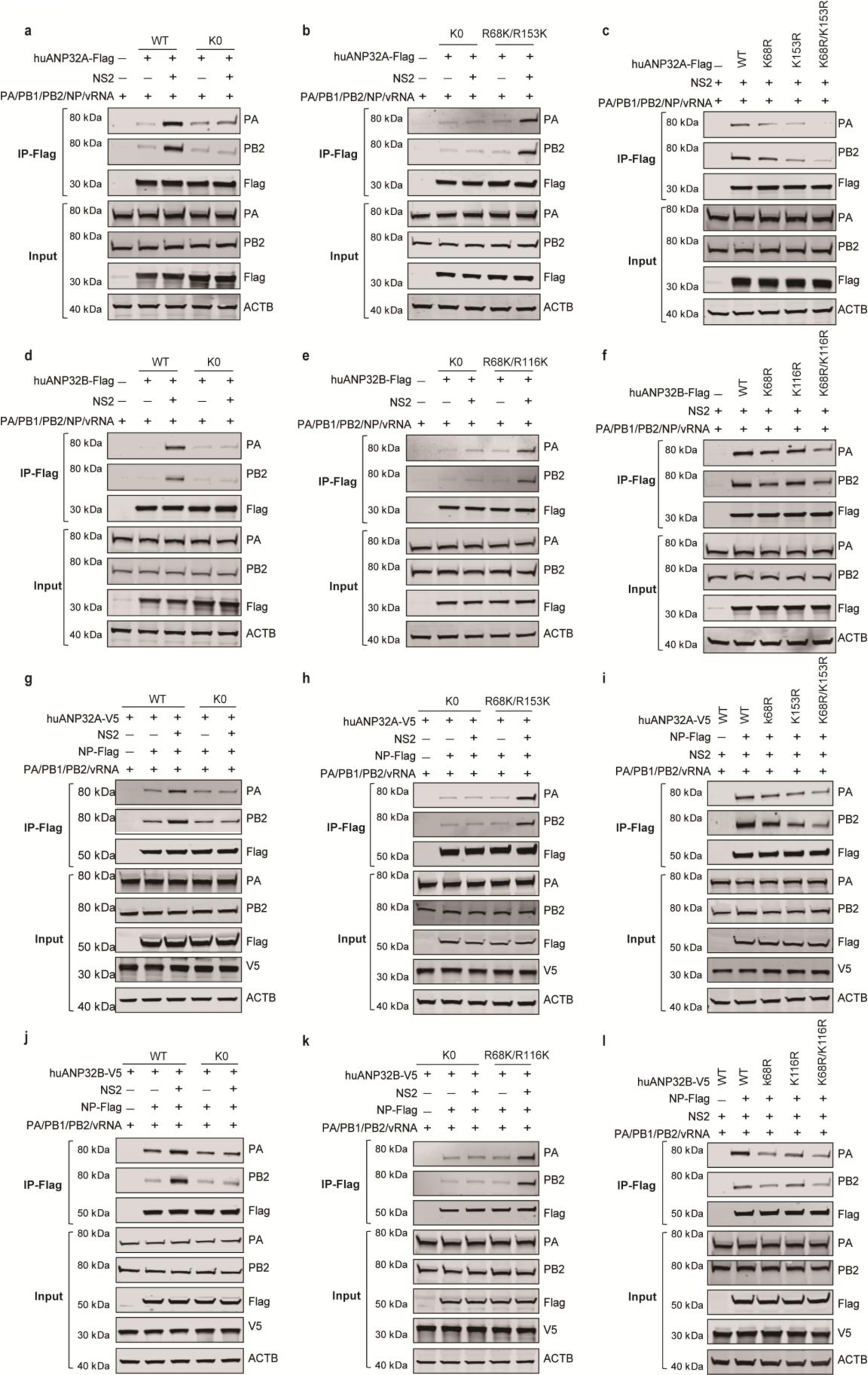
NS2-SIM promotes avian vPol activity by positively regulating AIV vRNP-huANP32A/B interactions as well as AIV vRNP assembly **a** NS2 failed to promote the binding of huANP32A-K0 to H9N2 vRNP. TKO cells were transfected with expression vectors for the indicated huANP32A-Flag constructs (0.4 μg), together with H9N2 PB1 (0.4 μg), H9N2 PB2 (0.4 μg), H9N2 PA (0.2 μg), H9N2 NP (0.8 μg), and vRNA-luciferase reporter (0.4 μg) and either with or without H9N2 NS2 (50 ng). The transfected cells were then subjected to immunoprecipitation and western blot analysis 24 h post-transfection. **b** NS2 promotes the binding of huANP32A-K0-R68K/R153K to H9N2 vRNP. **c** The effect of K68R, K153R or K68R/K153R mutations on the H9N2 vRNP-huANP32A interaction in the presence of NS2. **d** NS2 did not promote the binding of huANP32B-K0 to H9N2 vRNP. **e** NS2 promotes the binding of huANP32B-K0-R68K/R116K to H9N2 vRNP. **f** The effect of K68R, K116R or K68R/K116R mutations on the H9N2 vRNP-huANP32B interaction in the presence of NS2. For (b to f), experiments were performed as in Fig. 7a. **g** Comparison of the effect of NS2 on the H9N2 vRNP assembly in TKO cells reconstituted with either huANP32A or huANP32A-K0. TKO cells were transfected with different V5-tagged ANP32A (0.4 μg), Flag-tagged NP (0.8 μg) and polymerase plasmids from avian influenza viruses H9N2 (0.2 μg PA, 0.4 μg PB1, and 0.4 μg PB2) together with vRNA-luciferase reporter (0.4 μg) and H9N2-NS2 (50 ng). After anti-Flag precipitation at 24 h post-transfection, the indicated proteins were analyzed with western blotting. **h** Comparison of the effect of NS2 on the H9N2 vRNP assembly in TKO cells reconstituted with either huANP32A-K0 or huANP32A-K0-R68K/R153K. **i** Measurement of avian H9N2 vRNP assembly in HEK293T-TKO cells reconstituted with huANP32A or the indicated mutants. **j** Comparison of the effect of NS2 on avian vRNP assembly in TKO cells reconstituted with either huANP32B or huANP32B-K0. **k** Comparing the effect of NS2 on H9N2 vRNP assembly in TKO cells reconstituted with either huANP32B-K0 or huANP32B-K0-R68K/R116K. **l** Measurement of H9N2 vRNP assembly in HEK293T-TKO cells reconstituted with huANP32B or the mutants indicated. For (h to l), experiments were performed as in Fig. 7g. In (a to l), experiments were independently repeated three times with consistent results.

We proceeded to investigate the impact of huANP32A SUMOylation on the assembly of H9N2 vRNP in TKO cells when reconstituted with the indicated constructs. As shown in Fig. 7g, NS2 enhanced the H9N2 vRNP assembly in TKO cells when reconstituted with wild-type huANP32A, consistent with previous reports^22^. In contrast, NS2 failed to enhance the H9N2 vRNP formation in TKO cells reconstituted with huANP32A-K0 (Fig. 7g, h). However, when TKO cells were reconstituted with huANP32A-K0-R68K/R153K, we observed the enhancement of H9N2 vRNP assembly by the presence of NS2 (Fig. 7h).

Furthermore, in the presence of NS2, the level of H9N2 vRNP assembly was reduced when TKO cells were reconstituted with huANP32A-K68R/K153R, as compared to its wild type (Fig. 7i). For huANP32B, we performed similar experiments with the H9N2 vRNP and the indicated huANP32B constructs and obtained similar results (Fig. 7j-l).

Taken together, these data suggest that NS2 employs its SIM to interact with the SUMO of SUMOylated huANP32A/B, and this interaction promotes huANP32A/B-supported avian vPol activity and avian IAV replication by positively regulating AIV vRNP-huANP32A/B interactions as well as AIV vRNP assembly.

## Discussion

As critical cofactors for IAV vPol function, huANP32A/B efficiently support mammalian-adapted IAV vPol activity^11,13,14,27–30^. Nonetheless, huANP32A/B poorly support AIV vPol activity due to the lack of the specific 33 amino-acid insertion found in chANP32A, which serves as the molecular basis for the restriction of avian vPol activity in human cells^11–14,29,31,32^. AIVs have evolved various strategies to enable their polymerase to adapt to mammalian ANP32A/B. The predominant strategy used by most AIV strains is the acquisition of adaptive mutations. However, in recent work, we unveiled two previously unidentified strategies^22,33^. The first is that AIVs from avian hosts are able to package avian ANP32A to prime the early stage of viral replication in mammalian cells^33^. The second is that adaptation of the AIV polymerase to huANP32A/B is to some extent promoted with the help of the SIM in NS2^22^. However, the precise mechanism by which the SIM in NS2 promotes mammalian ANP32A/B-supported avian vPol activity remains elusive.

In this study, we continue to elucidate in more detail how NS2 employs its SIM to exert its AIV polymerase-enhancing function. Our findings reveal that NS2 uses its SIM to specifically interact with the SUMO in huANP32A/B. These SIM-SUMO interactions between NS2 and huANP32A/B are crucial for NS2 to promote avian vPol activity in human cells. We demonstrate that presence of SUMO at K68/K153 sites in huANP32A or K68/K116 sites in huANP32B dictates their binding affinity with NS2-SIM. Furthermore, disrupting the SIM-SUMO interactions between huANP32A/B and NS2, either by interfering with huANP32A/B SUMOylation or disrupting the integrity of the SIM inNS2, compromises the ability of NS2 to enhance avian vPol activity and AIV replication in human cells. This impairment stems from diminished interactions between AIV vRNP and huANP32A/B and reduced avian vRNP assembly.

SUMO modification is a highly conserved and important post-translational modification^16,17^, and plays a pivotal role in regulating diverse viral replication processes^34,35^. In the case of influenza viruses, several studies have shown that SUMOylation regulates viral replication levels and even the ability to spread at different stages of the viral replication cycle^36–40^. However, these investigations have predominantly centered on viral proteins themselves, and knowledge of the influence of SUMO modification of host factors on influenza virus replication remains relatively limited. As indispensable host factors for the function of influenza virus polymerase, huANP32A and huANP32B have been identified as potential SUMOylation substrates by large-scale proteomic analysis^23^. Here, we provide more evidence that huANP32A and huANP32B undergo SUMOylation, with neither showing a preference for SUMO1, SUMO2 or SUMO3. The SUMOylation process for huANP32A/B is orchestrated by the E3 SUMO ligase PIAS2α and the deSUMOylase SENP1. Interestingly, SUMOylation of huANP32A and huANP32B was up-regulated by H9N2 AIV infection. Our findings indicate that the SIM of NS2 recognizes and binds to SUMO in huANP32A/B. The expression of a NS2 mutant lacking a functional SIM, or huANP32A devoid of the SUMO acceptor K68/K153, or huANP32B lacking the SUMO acceptor K68/K116 all result in an impaired interaction between NS2 and huANP32A/B. Consequently, this reduction compromises the ability of NS2 to enhance huANP32A/B-supported AIV vPol activity. Collectively, our results underscore the indispensability of the SIM-SUMO interaction between NS2 and huANP32A/B for the role of NS2 in promoting avian vPol activity when supported by huANP32A/B.

Previous studies have demonstrated that huANP32A and huANP32B exhibit similar supportive roles in mammalian-adapted IAV vPol activity, and both show limited support for AIV vPol activity^12,13,32^. Despite our discovery in this study of the interaction between NS2 and huANP32A and huANP32B via a similar SIM-SUMO interaction pattern, there are distinctions between the two proteins. Specifically, huANP32A relies on SUMO at K68/K153 for its interaction with NS2-SIM, whereas huANP32B relies on SUMO at K68/K116. Intriguingly, sequence alignment analysis shows the conservation of K68/K116 across mammalian ANP32A/B, whereas K153 conservation is exclusive to mammalian ANP32A. While SUMOylation at the K116 site of huANP32A may also influence its interaction with NS2, our findings suggest that its significance is comparatively lower than that at the K68 and K153 sites. In addition, considering the substantial promotion of avian vPol activity by NS2 when supported by other mammalian ANP32A/B, we postulate that the functionality of these sites remains conserved across ANP32A/B from diverse mammalian species. Moreover, given the evolutionary conservation of these SUMOylation residues, we speculate that SUMOylation at these sites in huANP32A/B may also impact their function in supporting mammalian-adapted IAV vPol activity. Further research is necessary to validate this.

While we presented evidence highlighting the crucial role of the SIM-SUMO module in mediating the interaction between NS2 and huANP32A/B, we did not observe a complete disruption of this interaction upon mutation of the SUMO sites in huANP32A/B or the SIM site in NS2. This may be because, as other reports have shown^41^, rather than simply facilitating physical contact, SUMOylation may play a fundamental role in structurally orienting the two proteins for closer interaction. This hypothesis is further supported by the observation that both huANP32A-K68R/K153R and huANP32B-K68R/K116R retain comparable SUMOylation levels with their wild-type counterparts, yet still exhibit impaired interaction with NS2-SIM.

The molecular mechanisms underlying the maladaptation of AIV polymerase to huANP32A/B remain largely unknown. Studies using mammalian-adapted influenza virus polymerase subunits have shown that huANP32A/B functions in a manner dependent on its interaction with the trimeric polymerase complex^13,14^. However, some literature suggests that the interaction of huANP32A/B with the trimeric polymerase complex may not be species-specific^11^. The trimeric polymerase complex must associate with viral RNA and NP to form vRNPs, which then control transcription and replication of the viral genome^42^. A substantial body of research suggests that impaired assembly of avian vRNP is associated with the restriction of avian vPol activity in human cells^9,43,44^. Previously, we reported that the enhancement of avian vPol activity by NS2 in human cells occurs through promotion of the assembly of avian vRNP and facilitation of the interaction of vRNP with huANP32A/B^22^. In this study, we provide additional evidence linking the limitation of AIV vPol activity in human cells to compromised AIV vRNP assembly and attenuated interactions with huANP32A/B. We demonstrate that NS2 relies on SIM-SUMO interactions with huANP32A/B to surmount this restriction by fostering interactions between AIV vRNP and huANP32A/B and enhancing AIV vRNP assembly. However, the specific molecular mechanisms governing the role of ANP32A/B in supporting IAV vPol activity remain elusive. Notably, whether the enhanced assembly of avian vRNP precedes increased binding to ANP32A/B or vice versa remains uncertain. Nonetheless, studies have shown that ANP32A mediates the assembly of the influenza virus replicase by joining two polymerase molecules to form an asymmetric dimer, thus serving as an essential component of the influenza virus replicase^45,46^. This observation suggests the possibility that increased binding of the AIV vRNP to ANP32A/B may drive enhanced AIV vRNP assembly. Further investigation is required to confirm this.

The SIM typically comprises a central cluster of hydrophobic amino acids, often flanked by acidic or polar residues^19,20^. SIM motifs facilitate non-covalent interactions with mono- or poly-SUMOylated proteins. These SIM-SUMO interactions exhibit high instability, readily induced or suppressed by altering the SUMOylation status of the involved proteins. Such flexibility confers a survival advantage to cells, enabling swift assembly of protein complexes in response to changing stress conditions, as evidenced by numerous studies^21,47–49^. In this study, we offer additional evidence that, beyond cellular processes, invading viruses such as AIVs use the SIM-SUMO interaction paradigm to engage in host-virus interactions and finely regulate the assembly of protein complexes crucial for viral replication.

Based on our data, we propose a model elucidating the role of NS2 in promoting AIV vPol activity in human cells (Fig. 8). AIVs infection induces an elevation in huANP32A/B SUMOylation, subsequently recruiting NS2 through the SIM-SUMO interaction pattern to the nuclear replication platform, where viral genome replication takes place with the assistance of huANP32A/B. The recruited NS2 further enhances the functionality of huANP32A/B in supporting AIV vPol activity by facilitating the interaction of vRNP with huANP32A/B, thereby augmenting avian vRNP assembly. Given the dynamic and reversible nature of the SUMOylation process, the role of NS2 in promoting huANP32A/B-supported AIV vPol activity can be fine-tuned using this previously unidentified SIM-SUMO operating pattern. Our research contributes to comprehending the intricate interplay among NS2, huANP32A/B, and avian polymerase, shedding light on the molecular mechanism underlying the enhanced adaptation of avian vPol to huANP32A/B conferred by the SIM of NS2. Furthermore, our study delineates a SIM-SUMO-dependent mechanism through which the interaction between NS2 and huANP32A/B can be finely modulated by AIVs, allowing them to more effectively adapt to their new mammalian hosts.

**Fig. 8.**
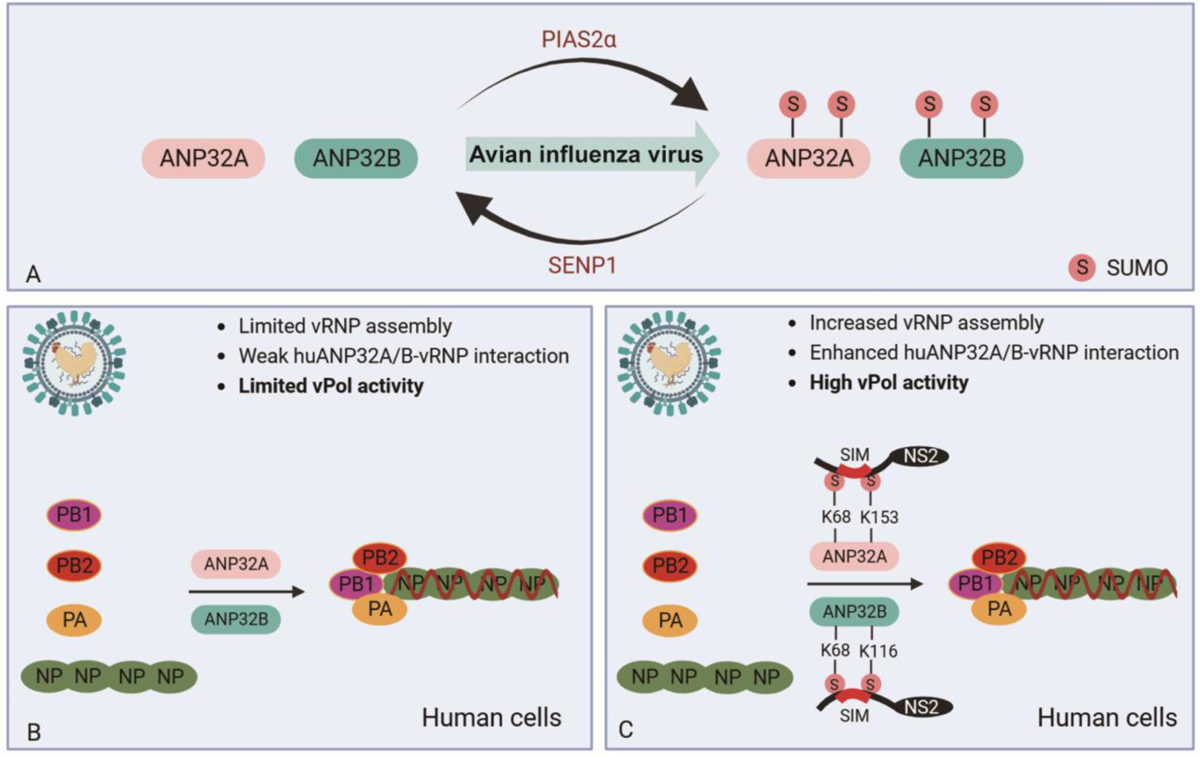
Working model for the role of NS2 in the promotion of huANP32A/B-supported AIV vPol activity. **a** huANP32A and huANP32B were SUMOylated by the E3 SUMO ligase PIAS2α, whereas they were deSUMOylated by SENP1. Both huANP32A and huANP32B SUMOylation increased upon AIVs infection. **b** huANP32A and huANP32B cannot efficiently support AIV vPol activity due to inefficient AIV vRNP assembly and weak AIV vRNP-huANP32A/B interactions. **c** During the AIVs infection process, the non-structural protein NS2 can be recruited to the replication platform by the SUMOylated huANP32A/B through the SIM-SUMO interaction pattern. Recruited NS2 uses its SIM to mediate intimate association with K68/K153-SUMO in huANP32A or K68/K116-SUMO in huANP32B, which in turn promotes huANP32A/B-supported AIV vPol activity by overcoming the defects in both AIV vRNP-huANP32A/B interactions and AIV vRNP assembly.

## Acknowledgements

We thank Dr. Zeijun Li (Shanghai Veterinary Research Institute, CAAS) for providing plasmids. We thank former lab members Zhenyu Zhang and Jiaqi Han for their help in the preparation of the MDCK-TKO cells. We thank former lab members Haili Zhang for providing plasmids. This work was supported by the National Natural Science Foundation of China (32330103) to X.J.W., the National Natural Science Foundation of China (32302959) to L.K.S., and the Natural Science Foundation of Heilongjiang Province of China (TD2022C006) to X.J.W.

## Author’s contributions

X.J.W. supervised the study and revised the manuscript. X.J.W. and L.K.S. designed the study and analyzed the data. L.K.S. wrote the original draft. L.K.S. performed the experiments. X.G. assisted with data validation. M.M.Y. and H.L.R. assisted with sample collection. X-F.W. assisted with manuscript editing. All authors reviewed the manuscript.

## RESOURCE AVAILABILITY

### Lead contact

Further information and requirements should be addressed to and will be fulfilled by lead contact Xiaojun Wang

### Materials availability

The materials used or generated in this study are available from the lead contact with a completed Materials Transfer Agreement.

### Data and code availability

- All data reported in this paper will be shared by the lead contact upon reasonable request.
- This paper does not report original code.
- Any additional information required to reanalyze the data presented in this work is available from the lead contact upon reasonable request.

## METHOD DETAILS

### Cells and plasmids

HEK293T cells, the previously described ANP32A/ANP32B/ANP32E triple knockout HEK293T cells (HEK293T-TKO)^24^, MDCK cells and the described ANP32A/ANP32B/ANP32E triple knockout MDCK cells (MDCK-TKO) were cultured in DMEM supplemented with 10% fetal bovine serum (Alphabio, Tianjin Alpha Biotechnology Co., Ltd) and penicillin/streptomycin (100 U/mL). Cells were grown in a 5 % CO_2_ cell culture incubator at 37°C.

The reverse genetic system for A/chicken/Zhejiang/B2013/2012 (H9N2) and plasmids pCAGGS-PB1, pCAGGS-PB2, pCAGGS-PA, pCAGGS-NP expressing the RNP components of H9N2 and H7N9 (PB2-627E) have been described previously^13,22,32^. The His-SUMO1/2/3 sequences and their non-conjugatable forms were subcloned seperately into a pCEF vector. Myc-tagged Ubc9 and V5/Flag-tagged ANP32A/B were subcloned into a pCAGGS vector. VN-myc-NS2, huANP32A/B-Flag-VC and GST-H9N2-NS2 were also subcloned into a pCAGGS vector. Flag-tagged SENP1, SENP2, SENP3, SENP5, SENP6, SENP7, USPL1, and Flag-tagged PIAS family members, including PIAS1, PIAS2α,

PIAS2β, PIAS3 and PIAS4 were subcloned seperately into a VR1012 vector. Gene sequences for the lysine-free mutant of huANP32A and huANP32B were generated by gene synthesis and inserted into the pCAGGS vector or VR1012 vector. The indicated additional mutations were introduced using a PCR method and were confirmed by DNA sequencing. For production of lentiviral particles, huANP32A/B and their different mutants were cloned into the lentivial vector pLVSIN-CMV-PGK-Puro for lentivirus production. All the constructs were subsequently verified by sequencing.

### Establishment of MDCK-TKO cells and stable cell lines

The canine ANP32A, ANP32B and ANP32E genes in MDCK cells were knocked out using the CRISPR-Cas9 system. Briefly, MDCK cells in 6-well plates were transfected with 1 μg of pMJ920 (Cas9-eGFP) plasmids and 1 μg of gRNA expression plasmids using PEI reagent. Single green fluorescent protein (GFP)-positive cells were harvested by using fluorescence-activated cell sorting (FACS) after transfection for 36 hours, and were further expanded for screening of monoclonal knockout cell lines with either western blotting or DNA sequencing. Single canine ANP32A knockout (AKO) and canine ANP32B knockout (BKO) cells were initially generated using two guides targeting the canine ANP32A locus and the canine ANP32B locus, respectively. Double canine ANP32A and ANP32B knock out (DKO) cells were then generated using either AKO cells with two guides against canine ANP32B locus or BKO cells with two guides against canine ANP32A locus. The final triple canine ANP32A, ANP32B and ANP32E knockout (TKO) cells were generated using DKO cells with two guides against canine ANP32E locus. AKO cells, BKO cells and DKO cells were validated with western blotting. TKO cells were validated with DNA sequencing. The guide RNAs used in this study are as follows: canine ANP32A (TAAGCGATAACAGAATCTCA and CACTTAAATTTAGATGCGTG), canine ANP32B (ATAGGTTCGAGAACTTGTCC and AGCCTACATTTATTAAACTG), canine ANP32E (TGACACACAGGCAATTATCA and TAATGTGGAACTAAGTTCAC).

For generation of MDCK-TKO cells stably reconstituted with the different Flag-tagged ANP32A/B constructs, different Flag-tagged ANP32A/B genes were first cloned into a pLVSIN-CMV-PGK-puro vector for producing lentiviral particles in HEK293T cells. MDCK-TKO cells were then infected with the generated lentiviral particles. Puromycin (1 μg/ml) was added 48 hours post-infection to obtain stable cell lines.

### Virus stock production, and infection assays

Avian H9N2 viruses were rescued in HEK293T cells by using the reverse genetic system of H9N2 and were further inoculated into SPF chicken embryos for propagation, as previously described^32^. The resulting viral stocks were then titrated in MDCK-chANP32A cells. For infection assays, plated cells were infected with H9N2 virus in Opti-MEM supplemented with 1 µg/ml TPCK. Cell supernatants were collected at the indicated time points, and virus titers were determined in MDCK-chANP32A cells.

### Minigenome assays

HEK293T-TKO cells plated in 24-well plates were co-transfected with the plasmids pPolI-luc (40 ng), pRL-TK encoding *Renilla* luciferase (5 ng), pCAGGS-PB1 (20 ng), pCAGGS-PB2 (20 ng), pCAGGS-PA (10 ng), pCAGGS-NP (40 ng), as well as plasmids encoding the indicated ANP32 proteins (20 ng) using PEI transfection reagent. The transfected cells were lysed 24 hours later after transfection, and the relative luciferase activity was measured using the Dual-Glo Luciferase Assay System (Promega) with a Berthold Centro LB 960 microplate luminometer.

### Immunoprecipitation assays

Cells were lysed with lysis buffer (50 mM Hepes-NaOH [pH 7.9], 100 mM NaCl, 50 mM KCl, 0.25% NP-40, and 1 mM DTT) plus complete inhibitor cocktail (APExBIO, Houston, USA, K1007). The lysates were centrifuged (12000 rpm, 10 min). The supernatants were incubated with anti-Flag M2 magnetic beads (Sigma-Aldrich, M8823) at 4°C overnight. The beads were washed five times with PBS and eluted with 3×Flag peptide (APExBIO, Houston, USA; A6001) and the immunoprecipitated proteins were then analyzed using western blotting.

### Mass spectrometry analysis

Samples were separated by SDS–PAGE and stained with Coomassie blue. Gel slices containing the proteins were collected for Mass spectrometry analysis as previously described^33^. All nano LC-MS/MS experiments were performed by J. Wang (Laboratory of Proteomics, Institute of Biophysics, Chinese Academy of Sciences, Beijing 100101, China). The mass spectrometry proteomics data have been deposited to the ProteomeXchange Consortium (https://proteomecentral.proteomexchange.org) via the iProX partner repository^50,51^ with the dataset identifier PXD053224.

### In vivo SUMOylation assays

Cells grown in 75 cm^2^ cell culture flasks were transfected with the indicated plasmids. Transfected cells were lysed in 2 mL RIPA lysis buffer supplemented with 1% SDS, 10 mM NEM (Sigma-Aldrich, E3876) and complete inhibitor cocktail (APExBIO, Houston, USA, K1007). The lysates were sonicated until they became fluid and then centrifuged (12000 rpm, 10 min). The supernatants were incubated with Ni2+-NTA beads (Sangon Biotech, C650033) at 4°C overnight. The beads were then washed sequentially with buffer A (50 mM Tris-HCl, 0.5 M NaCl, 6 M guanidium-HCl), buffer B (50 mM Tris-HCl, 0.5 M NaCl, 8 M urea), and PBS. The bound proteins were then eluted with elution buffer containing 200 mM imidazole, and analyzed with western blotting.

### siRNA treatment

siRNA transfection was performed using Lipofectamine™ 2000 transfection reagent according to the manufacturer’s protocol on the same day as HEK293T cells were seeded. 24 h after siRNA transfection, the cells were further transfected with plasmids as indicated using PEI transfection reagent. Cells were harvested for analysis 24 h after the second transfection step. siRNA oligonucleotides against SENP1 (siRNA1: 5’-GCGCCAGAUUGAAGAACAGAA-3’; siRNA2: 5’-UGACCAUUACACGCAAAGAUA-3’), PIAS2(siRNA1:5’-GCCAUGUUAUUACAGAGAUUA-3’;siRNA2:5’-GCUGCUAUUCCGCCUUCAUUA-3’) and non-targeting control (5’-UUCUCCGAACGUGUCACGU-3’) were purchased from Seven Innovation (China, Beijing) Biotechnology.

### Western blot analysis

Western blot analysis was conducted following established protocols^13^ using the following antibodies: rabbit anti-Flag (Sigma, F7425), rabbit anti-HA (Sigma, H6908), rabbit anti-ACTB (Abclonal, AC026), mouse anti-ACTB (Abclonal, AC004), rabbit anti-Myc (Abclonal, AE070), mouse anti-His (Proteintech, 66005-1-Ig), rabbit anti-V5 (Proteintech, 14440-1-AP), rabbit anti-SENP1 (Proteintech, 25349-1-AP), rabbit anti-GST (Proteintech, 10000-0-AP), rabbit anti-ANP32A (Proteintech, 15810-1-AP), rabbit anti-ANP32B (Proteintech, 10843-1-AP), rabbit anti-influenza A virus PB2 (GeneTex, GTX125926), mouse anti-influenza A virus PA (prepared in our laboratory, 1:5000 for WB), mouse anti-influenza A virus NP (prepared in our laboratory, 1:5000 for WB), Biotin-conjugated Affinipure Goat Anti-Rabbit IgG(H+L) (Proteintech, SA00004-2), DyLight™ 680-labelled, anti-mouse IgG (H + L) antibody (KPL, 072-06-18-06), DyLight™ 800-labelled, anti-rabbit IgG (H + L) antibody (KPL, 072–06-15-16) and DyLight™ 800-labelled streptavidin (KPL, 072-06-30-00).

To detect NS2-huANP32A/B interactions, the GST-NS2 signal within purified protein complexes was enhanced using the biotin-streptavidin system. Specifically, the membrane underwent sequential incubation with anti-GST (rabbit) antibody, Biotin-conjugated Affinipure goat anti-Rabbit IgG(H+L), and DyLight™ 800-labelled streptavidin. Subsequently, bands were analyzed by scanning blots with the Odyssey Imaging System (Li-Cor, Lincoln, NE, USA).

### Generation of stable cell lines

HIV-1-based lentiviral particles were produced by transfecting HEK293T cells with plasmids pspAX2, pMD2.0G and vector plasmids (pLVSIN-CMV-PGK-puro). Supernatants were collected 48 h post-transfection and filtered through 45 μM-pore size filters. Then MDCK-TKO cells were then infected with these generated lentiviruses. Stable cell lines were subjected to puromycin selection (1 μg/ml).

### Immunofluorescence staining

Immunofluorescence analysis was performed as previously described^52^. Briefly, the indicated cells were fixed in 4% paraformaldehyde for 20 min, followed by permeabilization in PBS containing 0.05% Triton-X-100 for 10 min. The cells were then blocked in PBS with 5% milk powder in PBS for 30 min before incubation with primary antibodies at 4°C overnight. The cells were then washed three times with PBS before incubation with secondary antibodies plus DAPI for 1 hour at room temperature. Images were obtained using a confocal microscope (Carl Zeiss LSM 800 Confocal Microscope and ZEN 2.3 LITE software).

### Flow cytometry

To detect BiFC signals, 293T cells were transfected with vectors expressing BiFC fusion proteins as indicated in Figures 1F and 1G. After 24 h, cells were analyzed using the fluorescein isothiocyanate (FITC) channel.

### RNA isolation and real-time quantitative PCR

Total RNA was isolated from cells using a RNeasy mini kit (Qiagen, 74106) according to the manufacturer’s instructions. Complementary DNA was synthesized using the PrimeScript RT reagent kit with a gDNA Eraser (Takara, RR047B) according to the manufacturer’s instructions. The mRNA level of PIAS2 in HEK293T cells were quantified using SYBR-Green (Takara, RR430A)-based real-time quantitative PCR analysis. Real-time PCR was performed using the human PIAS2 mRNA primers: PIAS2-qPCR-F: GTTCTTGGTGTCCAATGAGACCG; PIAS2-qPCR-R: TGCTTGCCTCACTGGCTACAGT. ACTB was used as a housekeeping control to normalize the number of living cells. The ACTB primers were ACTB-forward (5ʹACGGCATCGTCACCAACTG3ʹ) and ACTB-reverse (5ʹCAAACATGATCTGGGTCATCTTCTC3ʹ).

### Statistical analysis

All statistical analyses were performed with GraphPad Prism software. Quantitative data are presented as mean ± SD and statistical significance was analysed using unpaired Student’s t-tests, two-way ANOVA or one-way ANOVA followed by Dunnett’s multiple comparisons tests, where appropriate, as indicated in the figure legends. A p-value of 0.05 or less was considered statistically significant.

## Supplementary figures for

**Supplementary Fig. 1.**
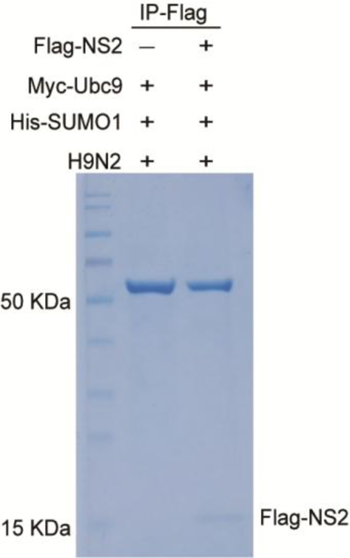
Identification of candidate proteins that associate with NS2 under the condition of H9N2 AIV infection HEK293T cells were transfected with the indicated plasmids, and 24 h later the transfected cells were infected with avian H9N2 virus (MOI = 0.01) for a further 24 h. The cells were then harvested for immunoprecipiation experiments using anti-Flag beads. The purified protein complexes were then analysed by SDS-PAGE followed by coomassie blue staining or using liquid chromatography-tandem mass spectrometry (LC-MS/MS) (see Table S1).

**Supplementary Fig. 2.**
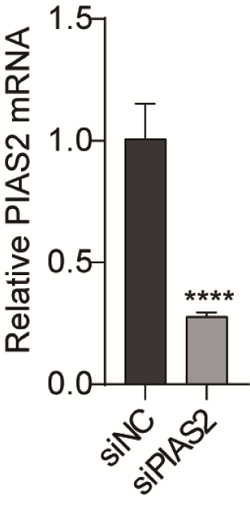
Treatment with siPIAS2 results in depletion of endogenous PIAS2 in HEK293T cells HEK293T cells were transfected with the indicated siRNA when the cells were plated in 6-well plates. After 48 hours, cells were harvested for RT-qPCR analysis. The 2^-ΔΔCt^ method was used to calculate PIAS2 mRNA expression relative to ACTB.

**Supplementary Fig. 3.**
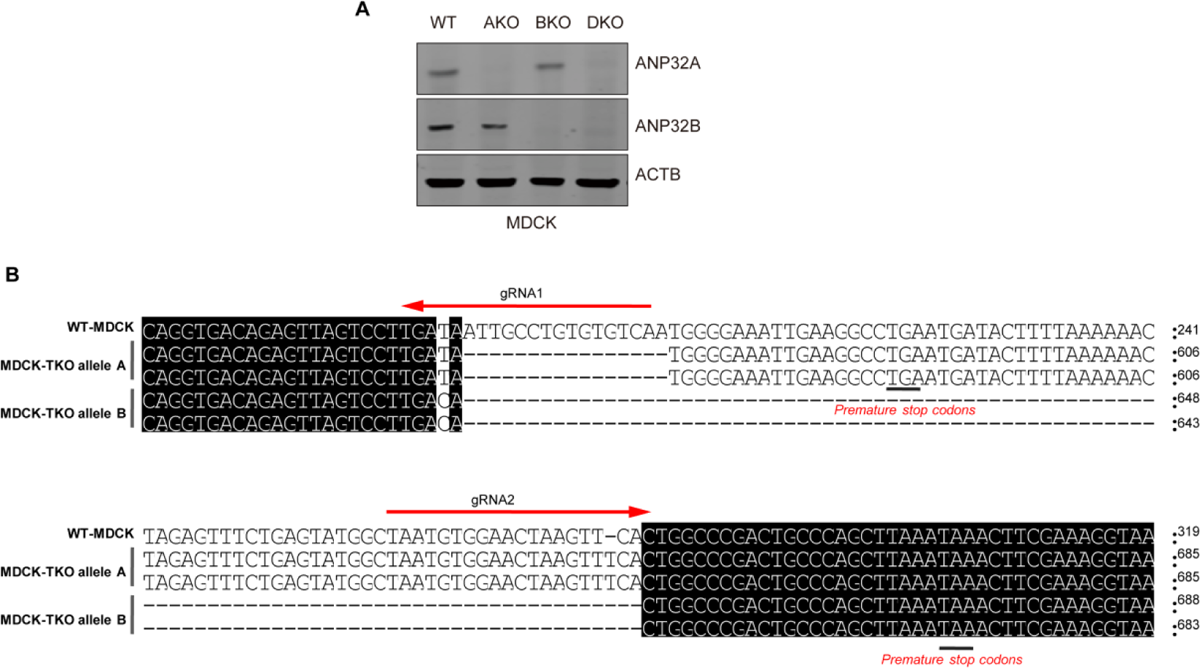
Establishment of the canine ANP32A, ANP32B and ANP32E triple knockout cell line (MDCK-TKO) a Western blot analysis of canine ANP32A and canine ANP32B protein in wild-type MDCK (WT), single canine ANP32A knockout cells (AKO), single canine ANP32B knockout cells (BKO) and double canine ANP32A and ANP32B knockout cells (DKO). b MDCK-TKO cells were generated from a DKO clone using CRISPR/Cas9 technology with two guides against the ANP32E locus. An independent TKO clone was verified by Sanger sequencing. Sequence alignment showed that both allele A and allele B had deletions resulting in premature stop codons in the ANP32E sequence.

**Supplementary Fig. 4.**
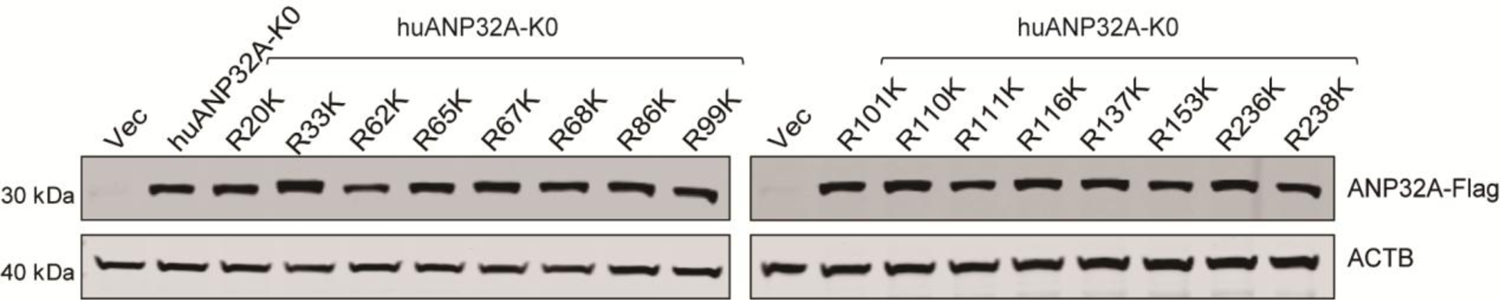
Immunoblotting analysis of lysates from HEK293T-TKO cells transfected with the indicated constructs HEK293T-TKO cells were transfected with the indicated plasmids for 24 hours. Cells were then harvested for immunoblotting analysis using specific antibodies.

**Supplementary Fig. 5.**
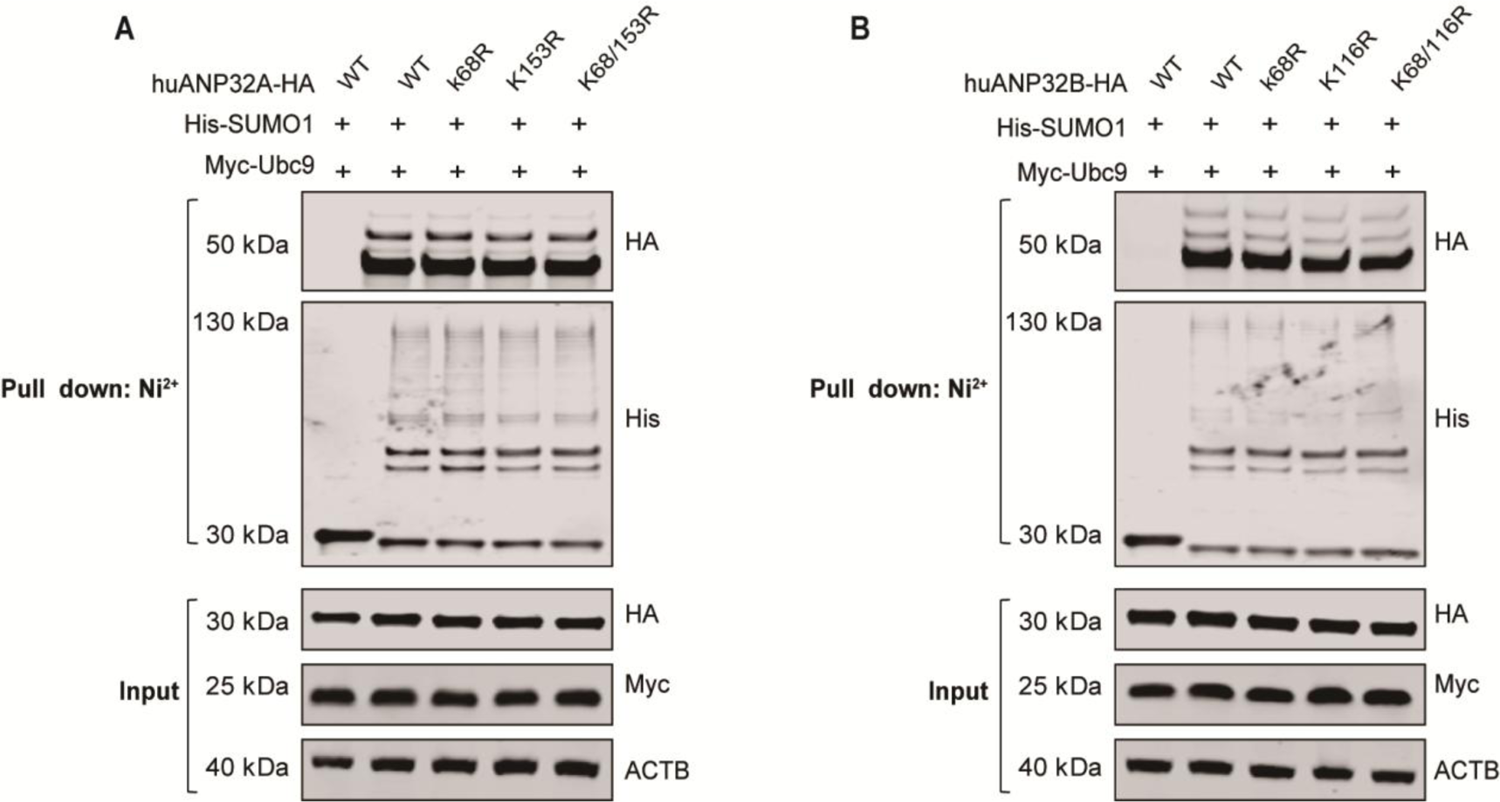
K68R/K153R mutation in huANP32A or K68R/K116R mutations in huANP32B did not affect their overall SUMOylation levels a HEK293T cells were transfected with His-SUMO1, and Myc-Ubc9, together with either huANP32A-HA or its indicated mutants for 24 hours, followed by in vivo SUMOylation assays and immunoblotting analysis. b HEK293T cells were transfected with His-SUMO1, and Myc-Ubc9, together with either huANP32B-HA or its indicated mutants for 24 hours followed by in vivo SUMOylation assays and immunoblotting analysis. In (a and b), experiments were independently repeated three times with consistent results.

**Supplementary Fig. 6.**
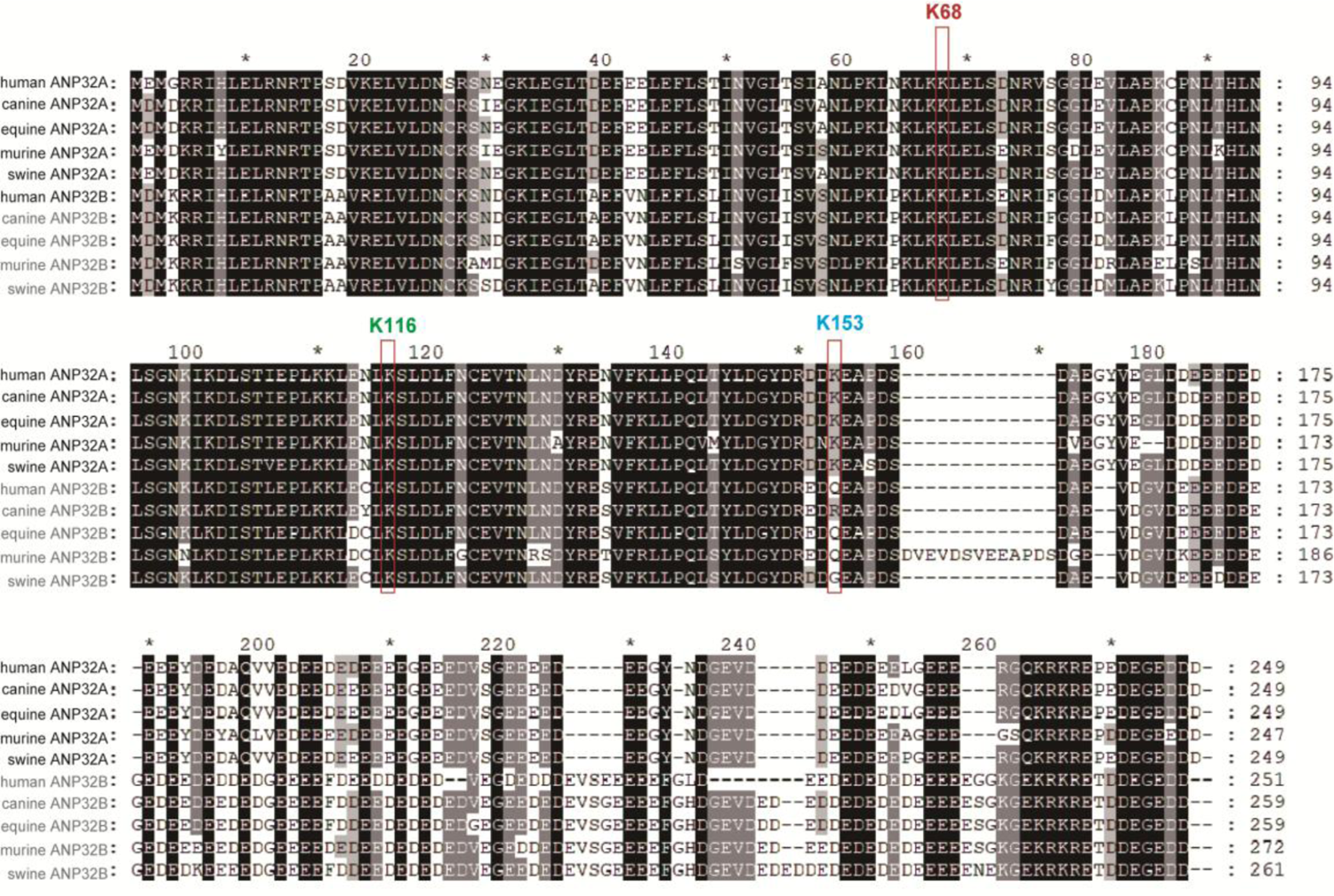
Sequence alignment of ANP32A and ANP32B proteins from different mammalian species ANP32A and ANP32B sequences from the indicated species were obtained from GenBank and included: human ANP32A (*Homo sapiens*, NP_006296.1); human ANP32B (*Homo sapiens*, NP_006392.1); swine ANP32A (*Sus scrofa*, XM_003121759.6); swine ANP32B (*Sus scrofa*, XM_021066477. 1); canine ANP32A (*Canis lupus familiaris*, NM_001003013.2); canine ANP32B (*Canis lupus dingo*, XM_025432382.3); equine ANP32A (*Equus caballus*, XP_001495860. 2); equine ANP32B (*Equus caballus*, XP_023485491.1); murine ANP32A (*Mus musculus*, NP_033802.2); murine ANP32B (*Mus musculus*, NP_570959.1). Sequence alignments were performed using DNASTAR Lasergene.

**Supplementary Fig. 7.**
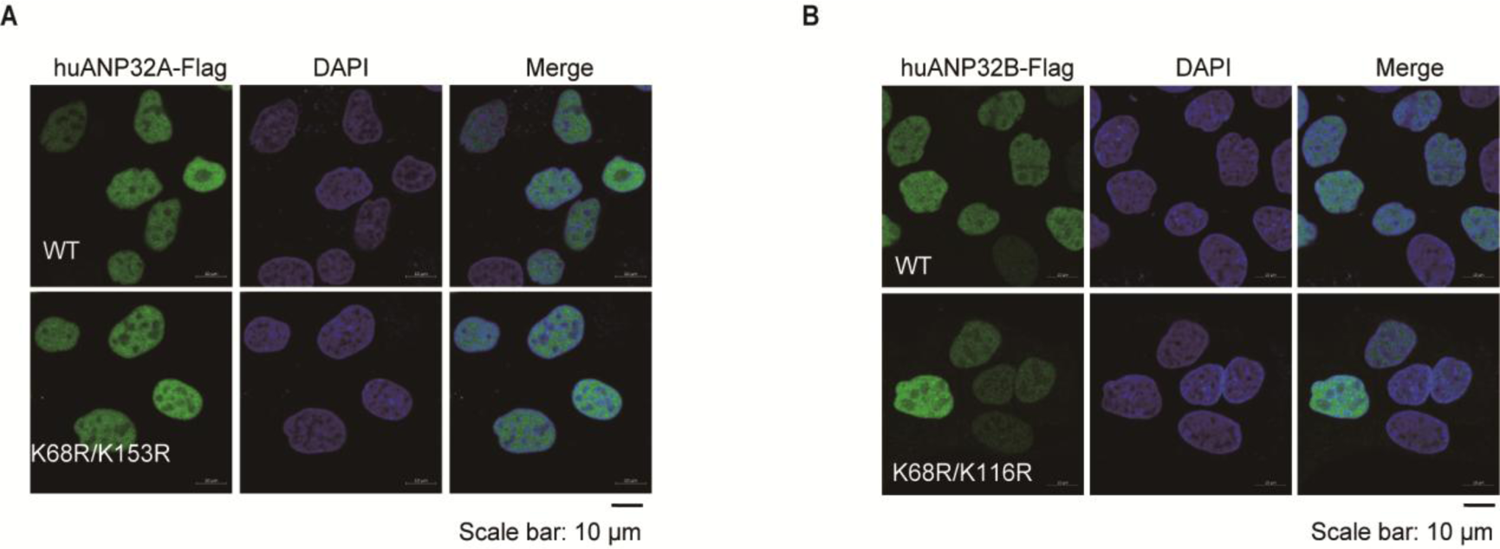
Neither K68R/K153R mutation in huANP32A nor K68R/K116R mutations in huANP32B affected their nuclear location, respectively a, b MDCK-TKO cells stably expressing huANP32A/B-Flag or their indicated mutants were subjected Immunofluorescence analysis.

**Supplementary Fig. 8.**
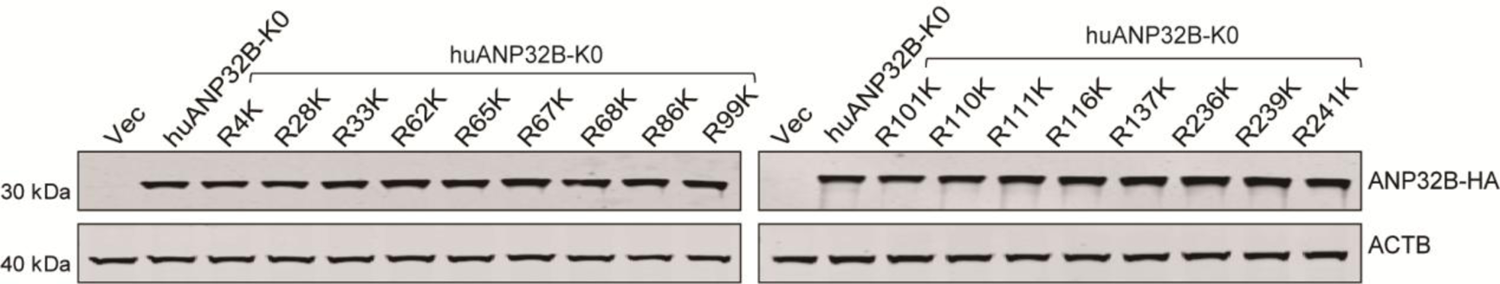
Immunoblot analysis of lysates from HEK293T-TKO cells transfected with the indicated constructs HEK293T-TKO cells were transfected with the indicated plasmids for 24 hours. Cells were then harvested for immunoblotting analysis using specific antibodies.

